# The effects of phenylalanine and tyrosine levels on dopamine production in rat PC12 cells. Implications for treatment of phenylketonuria and tyrosinemia type 1

**DOI:** 10.1101/2023.09.05.556372

**Authors:** Peter D. Szigetvari, Sudarshan Patil, Even Birkeland, Rune Kleppe, Jan Haavik

## Abstract

Phenylketonuria (PKU) is an autosomal recessive metabolic disorder caused by mutations in the phenylalanine hydroxylase (PAH) gene, resulting in phenylalanine accumulation and impaired tyrosine production. In Tyrosinemia type 1 (TYRSN1) mutations affect fumarylacetoacetate hydrolase, leading to accumulation of toxic intermediates of tyrosine catabolism. Treatment of TYRSN1 with nitisinone results in extreme tissue levels of tyrosine. Although PKU and TYRSN1 have opposite effects on tyrosine levels, both conditions have been associated with neuro- psychiatric symptoms indicative of impaired dopamine (DA) synthesis. However, concrete *in vivo* data on the possible molecular basis for disrupted DA production under disease mimicking conditions have been lacking. In pursuit to uncover associated molecular mechanisms, we exposed an established, DA producing cell line (PC12) to different concentrations of phenylalanine and tyrosine in culture media. We measured the effects on viability, proteomic composition, tyrosine, DA and tyrosine hydroxylase (TH) levels and TH phosphorylation. TH catalyzes the rate-limiting step in DA synthesis. High extracellular levels of phenylalanine rapidly depleted cells of intracellular tyrosine and DA. Compared to physiological levels (75 µM), either low (35 µM) or high concentrations of tyrosine (275 or 835 µM) decreased cellular DA, TH protein, and its phosphorylation levels. Using deep proteomic analysis, we identified multiple proteins, biological processes and pathways that were altered, including enzymes and transporters involved in amino acid metabolism. Specifically, levels of the broad specificity transporter of neutral amino acids LAT1 and the branched-chain amino-acid-transporter BCAT2 were both similarly increased. Using this information and published data, we developed a mathematical model to predict how extracellular levels of aromatic amino acids can affect the cellular synthesis of DA *via* different mechanisms. Together, these data provide new information about the normal regulation of neurotransmitter synthesis and how this may be altered on the molecular level in neurometabolic disorders, such as PKU and TYRSN1.

**Highlights:** Tyrosinemia type I and phenylketonuria are rare metabolic disorders, characterized by extremely elevated plasma levels of the aromatic amino acids tyrosine and phenylalanine, respectively. To study the molecular consequences of dysregulated amino acid uptake on dopamine homeostasis, we simulated these conditions using rat pheochromocytoma (PC12) cells, an established model system for investigating catecholamine production. Non-physiological tyrosine levels induced a broad cellular response, including reduced protein level and phosphorylation stoichiometry for tyrosine hydroxylase, the rate-limiting enzyme in catecholamine biosynthesis, which in turn resulted in decreases in both intra- and extracellular dopamine levels. Similarly, dopamine content was also decreased when cells were exposed to high phenylalanine levels characteristic of phenylketonuria, although the underlying molecular mechanism for this response appears to differ.

## 1 | Introduction

Neurometabolic diseases are typically caused by genetic variants affecting critical protein function, thus interfering with the normal development and activity of the nervous system. In particular, metabolic disorders affecting catecholaminergic neurotransmission have been linked to symptoms and aetiologies of several neuropsychiatric disorders (Cannon Homaei et al., 2022). Such inborn errors of metabolism that also affect the nervous system include the aminoacidopathies phenylketonuria (PKU; OMIM 261600) and tyrosinemia type 1 (TYRSN1; OMIM 276700; (Pohorecka et al., 2012)). Both PKU and TYRSN1 share some symptoms with neuropsychiatric disorders, notably attention deficit hyperactivity disorder (ADHD, OMIM 143465) (Pohorecka et al., 2012; Stevenson & McNaughton, 2013), and have been associated with low prefrontal dopamine levels (Antshel & Waisbren, 2003; Borodovitsyna et al., 2017; Volkow et al., 2009).

PKU is an inborn error of metabolism resulting in decreased metabolism of phenylalanine (L-Phe), most commonly caused by mutations in *PAH.* TYRSN1 is an autosomal recessive disorder caused by loss of function mutations in the fumarylacetoacetate hydrolase (*FAH*; EC 3.7.1.2) gene, which encodes the terminal enzyme within the tyrosine (L-Tyr) catabolic pathway. The resulting accumulation of toxic metabolites such as fumarylacetoacetate and succinylacetone is associated with a severe, life-threatening condition, leading to multiple organ failure involving the liver and kidneys, causing cirrhosis or hepatocellular carcinoma and death if left untreated (Mayorandan et al., 2014). Although the estimated worldwide incidence of the disease is low (1:100 000 – 120 000 live births), certain geographical and ethnic areas of higher occurrence have also been identified (Bliksrud et al., 2012; De Braekeleer & Larochellet, 1990).

Nitisinone (2-(2-nitro-4-trifluoromethylbenzoyl)-1,3-cyclohexanedione, (NTBC)) prevents the formation of carcinogenic metabolites *via* the inhibition of 4-Hydroxyphenylpyruvate dioxygenase (HPPD), effectively halting tyrosine degradation (Lindstedt et al., 1992). When treatment is initiated early, NTBC dramatically improves prognosis (Spiekerkoetter et al., 2021), however, tissue tyrosine may reach extremely high levels, even with adherence to a phenylalanine and tyrosine restricted diet (Bendadi et al., 2014). Neurocognitive deficits are prominent among treated TYRSN1 patients. Reported problems include learning difficulties (Masurel-Paulet et al., 2008), attention deficit (Pohorecka et al., 2012), difficulties with social cognition (van Vliet et al., 2019) and working memory (Van Ginkel et al., 2016), lower IQ, as well as suboptimal motor function (Cannon Homaei et al., 2022; Thimm et al., 2012). It has been reported that underperformance on cognitive assessments reflects the L-Tyr concentrations measured in plasma, where patients with the highest L-Tyr levels also show most severe neurocognitive deficits (Barone et al., 2020; Walker et al., 2018). Several explanations have been offered for the observed neurodevelopmental problems, including a direct toxic effect of high tissue L-Tyr, pre-treatment liver dysfunction, imbalance of the influx of amino acids through the BBB, decreased serotonin in the CNS, or direct side-effects of the NTBC treatment itself (van Ginkel et al., 2017).

As tyrosine is the preferred substrate for tyrosine hydroxylase (TH, EC 1.14.16.2), the rate-limiting enzyme in the biosynthesis of dopamine (and other catecholamines) (Nagatsu et al., 1964), it has been suggested that the observed cognitive symptoms are related to the pathologically elevated substrate levels for TH. Consequently, this was speculated to increase tyrosine hydroxylation, thus elevating dopamine levels in the human brain (van Ginkel et al., 2017), although, homovanillic acid (HVA) measurements in the cerebrospinal fluid (CSF) of patients were not able to establish this possible outcome (Thimm et al., 2011).

Contrasting with the proposed increase in dopamine levels in NTBC treated TYRSN1, the overlapping cognitive deficits observed in PKU and ADHD were suggested to result from decreased catecholaminergic neurotransmission (Antshel & Waisbren, 2003; Harding et al., 2014) (Borodovitsyna et al., 2017; Volkow et al., 2009). It was postulated that hypotyrosinemia is typical in PKU (Hanley et al., 2000) and that pathologically elevated L-Phe levels may inhibit brain production of both serotonin and dopamine in PKU patients (Lykkelund et al., 1988), however, the exact molecular mechanisms remain unclear. Intriguingly, the cognitive symptoms in PKU, ADHD and TYRSN1 all respond to treatment with centrally acting stimulant drugs that increase synaptic levels of dopamine and noradrenaline (Arnold et al., 2004; Barone et al., 2020).

The biosynthesis of dopamine has been intensively investigated, mainly focusing on the regulation of TH protein turnover and the specific activity of this enzyme. Long-term regulation of TH protein abundance occurs at the transcriptional level and possibly through ubiquitin-mediated protein degradation (Døskeland & Flatmark, 2002; Kawahata & Fukunaga, 2020; Nakashima et al., 2018), while short term regulation of TH activity relies on the reversible phosphorylation at N-terminal serines 8, 19, 31 & 40 (Ghorbani et al., 2020; Kaufman, 1995) as well as end product inhibition by L-DOPA and catecholamines, including DA (Bongiovanni et al., 2006; Kaufman, 1995). In addition to end-product inhibition, TH is subject substrate inhibition by tyrosine. This inhibition by excess tyrosine (>50 µM) already occurs at physiological levels in humans (DePietro & Fernstrom, 1998; Ribeiro et al., 1991) and is believed to be important to stabilize the rate of DA synthesis, irrespective of dietary fluctuations in tyrosine availability (Best et al., 2009; Reed et al., 2010). Taking this into consideration, we proposed that in respect to cognitive symptoms, the possibility of a shared pathophysiological mechanism between PKU, ADHD and TYRSN1 could not be excluded. We argued that that in humans, abnormally elevated L-Tyr concentrations - paradoxically - may exert attenuation on catecholamine signaling in the PFC via the mechanism of substrate inhibition on TH. This would likely affect serotonin biosynthesis as well through competitive inhibition of tryptophan hydroxylase 2 (TPH2) and lowered tryptophan (L-Trp) availability (Barone et al., 2020).

As in humans, treatment of mice with NTBC increases tissue levels of tyrosine (van Ginkel et al., 2022). However, unlike in humans, where elevated plasma levels of tyrosine were associated with impaired cognitive functioning (Barone et al., 2020), no alterations in either behavior or brain monoamine levels were observed in this mouse model. This indicates that there is no simple relationship between tissue levels of tyrosine and brain dopamine and that the synthesis of this transmitter is tightly regulated. Indeed, mathematical modeling suggests a remarkable resilience to sudden changes in environmental precursor concentrations that fall within physiological parameters (Best et al., 2009; Reed et al., 2010). Still, due to the conflicting results reported, and particularly the apparent discrepancies between *in vitro* and *in vivo* studies, many questions remain regarding the aetiology of cognitive deficits in treated TYRSN1. This is partially due to limitations of available experimental approaches. In humans, measurements of tyrosine and dopamine levels are limited to blood and cerebrospinal fluid (CSF), while in animal models, tissue levels can also be measured. However, as brain tissue consists of hundreds of cell types and the relevant enzymes, transporters and metabolites are highly compartmentalized, studies of bulk tissue extracts provide limited mechanistic insight.

Here we investigated the effects of acute and chronic alterations in extracellular levels of tyrosine and phenylalanine on catecholamine synthesis and amino acid homeostasis in rat phaeochromocytoma (PC12) cells. We show that exposure to both pathologically low or high levels of tyrosine reduces cellular dopamine, as well as TH protein and phosphorylation levels. Similarly, proteomic analyses show that several biological processes relevant to the metabolism of tyrosine and related amino acids are altered at non-physiological tyrosine concentrations. Such mechanisms may explain why high levels of circulating tyrosine may induce a dopamine deficient state in the CNS. In contrast, elevated phenylalanine levels appear to reduce dopamine production primarily through restricted tyrosine uptake by the affected cells.

## 2 | Methods

### 2.1 | Cell culture

Chemicals used in this study were purchased from Merck KGaA, Germany unless otherwise stated. A rat pheochromocytoma (PC12) cells sub-clone (Sannerud et al., 2006) was kindly provided by Ann Kari Grindheim (UiB, Department of Biomedicine, Bergen, Norway), and maintained in T75 TC-flasks (REF 83.3911.002; Sarstedt AG, Germany) in an incubator set at 37°C with an atmosphere of 5% CO_2_ and 95% humidity (Greene et al., 1987). Cells were fed every third day and passaged weekly when reaching ∼70% confluency. The cell culture medium was composed of Ham’s F-10 GlutaMAX™ Nutrient Mix (REF 41550-021; Gibco/Thermo Fischer Scientific Inc.) containing 10% heat-inactivated horse serum (#26050088, ThermoFisher) and 5% fetal bovine serum (#10100147, ThermoFisher), as well as 50 U/mL of penicillin/streptomycin. Cells grew as expected with normal yields using this growth medium. Although PC12 cells are most frequently grown in RPMI media, for our purposes, we required a nutrient mixture with very low L-Tyrosine and L-Phenylalanine content, that with the addition of both horse serum as well as FBS only rose to about 35 and 40 µM, respectively (Elhassan et al., 2001; Westfall et al., 1954). Thus, the final amino acid composition of the F10 media was as follows (in µM): Glycine (129.9); L-Alanine (163.4); L-Alanyl-L-Glutamine (1000.0); L-Arginine hydrochloride 1017.5); L-Asparagine-H2O (102.6); L-Aspartic acid (105.9); L-Cysteine (218.0); L-Glutamic acid (142.3); L-Histidine hydrochloride-H2O (123.9); L-Isoleucine (51.5); L-Leucine (131.3); L-Lysine hydrochloride (187.2); L-Methionine (36.5); L-Phenylalanine (39.2); L-Proline (110.1); L-Serine (116.2); L-Threonine (61.5); L-Tryptophan (13.6); L-Valine (57.8); L-Tyrosine (35.2). The only difference between growth medium and treatment medium is that in the latter, L-Tyrosine concentration was left as is or raised to 75, 275 & 835 µM, respectively. In separate experiments, L-Tyr concentration was increased to 75 µM, while L-Phe levels were adjusted to 100, 250, 500 & 1000 µM, respectively. For experiments, the 4^th^ passage was used of each thawed batch of cells in order to avoid inconsistencies.

### 2.2 | Treatment conditions

Cells were plated on Nunclon™ Delta Surface flat-bottomed 24-well plates (#142475; Thermo Fisher Scientific Inc.), each well containing 1.0 x10^5^ cells in 1 ml media as described above. Then, the cultures were grown for 48 hours prior to the start of the experiment in fresh media containing basal concentrations of amino acids present within the complete nutrient mixtures. Treatment with media containing L-Tyr at concentrations corresponding to hypotyrosinemic (35 µM), physiological (75 µM), and conditions typical at the onset of mild (275 µM) and more severe (835 µM) cognitive symptoms observed in treated TYRSN1 were conducted over periods of 1, 3, 6 & 24 hours. Separate experiments with L-Phe-containing media mimicking conditions of hypophenylalaninemia (40 µM), physiological (100 µM), mild or asymptomatic PKU (250 µM) and PKU displaying cognitive deficits (500 & 1000 µM) lasted 1, 3 & 6 hours. Once the incubation periods concluded, dishes containing treated cells destined for monoamine measurements were placed on ice, washed twice with Dulbecco’s phosphate-buffered saline (Gibco), and harvested by adding 150 µl chilled lysis buffer composed of 2N HClO_4_, 2.5 mM EDTA, 5 mM sodium metabisulphite. The bottom surface of the wells were then scraped, and the resulting suspensions were collected in 1.5 ml Eppendorf tubes. Samples obtained this way were centrifuged at 16000 RCF for 15 min at 4°C. Finally, supernatants were stored at −80°C until analysis. For protein determination using western blotting and BCA protein assay (Pierce, #23227), cells were treated in parallel (identically from the fourth passage), but instead of 2N HClO_4_, cells were harvested using Pierce™ IP Lysis buffer (# 87787, Thermo Fisher Scientific Inc.).

CellTiter-Blue (Promega #G808B) test using the fluorimetric resazurin reduction technique was carried out to estimate the number of viable cells in response to the described treatments (Waløen et al., 2021). Liquid nitrogen stocks of PC12 cells were plated on Falcon flat-bottomed 96-well plates) (#353219; BD Falcon), each well containing ∼3.0 10^4^ cells in 100 µl growth media. Then, cells were grown for 24 hours prior to the start of the experiment with media containing basal concentrations of amino acids present within the complete nutrient mixtures, as described above. Treatments were conducted over periods of 1, 6, 18, 24,36 & 48 hours with fresh treatment media containing L-Tyr concentrations of 35, 75, 275 & 835 µM. After injecting 20 µL of CellTiter Blue reagent to each well, the plates were incubated for 2 hours. Fluorescence (λ=590) was measured with a Victor 3 1420 Multilabel plate reader. Light microscopic observations presented no evidence for cellular stress regardless of the L-Tyr concentration or the length of the treatment.

### 2.3 | Monoamine analysis

Cell lysate samples were thawed on ice and filtered prior to analysis. DA content in lysates was measured using high-performance liquid chromatography (HPLC) equipped with fluorometric detection (Agilent) and a Zorbax Eclipse Plus C18 column (125 × 4.6 mm; 3.5 µm pore size (Agilent)). The mobile phase was composed of 13% methanol, 3% acetonitrile, 1% 50 mM Na-acetate pH 4.0 and 83% 75 mM Na- dihydrogen phosphate pH 3.0; 1.7 mM 1-octanesulfonic acid sodium salt, 25 µM Na-EDTA; 0.01 %V/V triethylamine. The flowrate was set to 1.0 ml/min at 30°C with 20 min run-time. Wavelengths for ε_x_ & ε_m_ were 280 and 330 nm, respectively. L-Tyr and dopamine eluted at ∼3.45 and ∼7.6 min, respectively (Fig. S1). Peak integrations and peak area quantifications were carried out using Agilent’s ChemStation software. For statistical comparisons, one-way ANOVA with Dunnett’s multiple comparisons tests were performed using GraphPad Prism 9 (version 9.5.1; San Diego, CA). Unless otherwise specified, all data are expressed as mean ± SD.

### 2.4 | SDS–PAGE and immunoblotting

PC-12 cells were treated with 35, 55, 75, 115, 275 & 835 µM of L-tyrosine dissolved in complete F10 cell culture medium for 1 & 6 hrs. and lysed in lysis buffer containing 50 mM Tris, 100 mM NaCl, 1 mM EDTA, NP-40 0.5%, 1 mM dithiothreitol, 1 mM Na_3_VO_4_, 50 mM NaF, and 1× protease inhibitor cocktail from Roche #11836170001. The homogenate was centrifuged for 10 min at 14000 x g at 4°C. Protein concentration was measured using the BCA protein assay (Pierce, #23227). Homogenates were stored at −80°C until use. About 15 μg of protein samples from lysates were boiled at 95°C for 5 min in Laemmli sample buffer (Bio-Rad) and resolved in 4-15% gradient SDS-PAGE gels (Mini-Protean TGX; Bio-Rad) and the proteins were transferred onto nitrocellulose membranes (0.2 µm pore size) by blotting performed using the Trans-Blot Turbo Transfer System (Bio-Rad; Hercules, USA) essentially according to the manufacturer (25V/1,3 mA, 10 min transfer). The membranes were blocked with 5% BSA and probed against P-^S19^-TH (#PA5-104765, ThermoFisher Scientific; 1:1000), P-^S40^-TH (#p1580-40, Phosphosolutions; 1:1000), TH (#66334-1-IG, Proteintech; 1:1000), Pan14-3-3 (#sc-629, Santa Cruz Biotech; 1:1000), DHFR (#133546, Abcam; 1:1000), QDPR (#186411, Abcam; 1:1000), BCAT2 (#95976, Abcam; 1:1000), GAPDH (#sc-32233 Santa Cruz Biotech; 1:1000) primary antibodies and proteins were detected by incubation at a 1:5000 dilution with secondary horseradish peroxidase (HRP)- conjugated antibodies (#170-6516, goat anti-mouse and 170-6515; goat anti-rabbit from Bio-Rad, Hercules, USA). The reactive protein bands were visualized using the WesternBright ECL HRP substrate (Advansta; San Jose, USA). The blots were scanned using Gel DOC XRS+ (Bio-Rad) and densitometric analyses were performed with Image J software (NIH, Bethesda, MD). Welch’s T-test was applied to determine significance against physiological values (values are expressed as mean ± SD).

### 2.5 | Sample Preparation (SP3) and HPLC tandem mass spectrometry (MS) Analysis

PC-12 cells were treated with 35, 75, 275 & 835 µM of L-tyrosine-containing media for 6 hours as described above and lysed in Pierce™ RIPA lysis buffer. Disulfide bonds were reduced using 5mM (2- carboxyethyl) phosphine (TCEP). The reduced cysteine residues were alkylated using 10mM chloroacetamide to prevent reformation of the disulfide bonds. SP3 reaction: the samples were mixed with paramagnetic beads (Sera-Mag Speed beads, GE healthcare) in a 1:10 ratio and subjected to agitation, before the addition of 100% ethanol to 70% ethanol final concentration. This solution was agitated at 1000 rpm for 7 min at RT. Magnetic separation: the SP3 reaction mixture was placed in a magnetic field, and the supernatant was removed. This procedure was repeated twice with 80% ethanol to remove any remaining lysis-buffer. Elution: the bound peptides were eluted from the beads using a high salt buffer (0.5M NaCl) and subsequently desalted using Oasis spin columns (Waters, MA, USA).

About 0.5 μg protein as tryptic peptides dissolved in 2% acetonitrile (ACN) and 0.5% formic acid (FA) were injected into an Ultimate 3000 RSLC system (Thermo Scientific, Sunnyvale, CA, USA) connected online to Orbitrap Eclipse mass spectrometer (Thermo Scientific) equipped with EASY-spray nano-electrospray ion source (Thermo Scientific). For trapping and desalting processes, the samples were loaded and desalted on a pre-column (Acclaim PepMap 100, 2 cm × 75 µm ID nanoViper column, packed with 3µm C18 beads) at a flow rate of 5 µL/min for 5 min with 0.1% TFA (trifluoroacetic acid). Peptides were separated during a biphasic ACN gradient from two nanoflow UPLC pumps (flow rate of 250 nL/min) on a 25 cm analytical column (PepMap RSLC, 50 cm × 75 µm ID EASY-Spray column, packed with 2 µm C18 beads). Solvent A and B were 0.1% FA (vol/vol) in dH2O and 100% ACN respectively. The gradient composition was 5% B during trapping (five minutes) followed by 5–7% B over 30 second, 8–22% B for the next 145 min, 22–28% B over 16 min, and 35–80% B over 15 min. Elution of very hydrophobic peptides and conditioning of the column were performed during 15 min isocratic elution with 90% B and 20 min isocratic elution with 5% B respectively. The eluting peptides from the LC-column were ionized in the electrospray and analyzed by the Orbitrap Eclipse. The mass spectrometer was operated in the DDA-mode (data-dependent-acquisition) to automatically switch between full scan MS and MS/MS acquisition. Instrument control was achieved through Tune 2.7.0 and Xcalibur 4.4.16.14.

Survey full scan MS spectra (from m/z 375 to 1500) were acquired in the Orbitrap with resolution R = 120,000 at m/z 200 after accumulation to a target value of 4e5 in the C-trap, ion accumulation time was set as auto. FAIMS was enabled using two compensation voltages (CVs), −45V and −65V respectively. During each CV, the mass spectrometer was operated in the DDA-mode (data-dependent-acquisition) to automatically switch between full scan MS and MS/MS acquisition. The cycle time was maintained at 0.9s/CV. The most intense eluting peptides with charge states 2 to 6 were sequentially isolated to a target value (AGC) of 2e5 and maximum IT of 120 ms in the C-trap, and isolation width maintained at 0.7 m/z, before fragmentation was performed with a normalized collision energy (NCE) of 30%, and fragments were detected in the Orbitrap at a resolution of 30 000 at m/z 200, with first mass fixed at m/z 110. The spray and ion-source parameters were as follows: ion spray voltage = 1900 V, no sheath and auxiliary gas flow, and capillary temperature of 275 °C.

### 2.6 | Statistical and bioinformatic analysis

The LC-MS raw files were searched in Proteome Discoverer version 2.4 (ThermoFisher). For GO analysis, filtered gene lists split to highlight genes differentially upregulated or downregulated in each dataset were individually submitted to DAVID and GO annotation gathered for Biological Function, Molecular Function and Cellular Component. GO terms with FDR < 0.05 were considered significantly enriched. Protein-protein interaction analysis for the differentially expressed proteins was performed by using the online tool STRING (version 11.5). This was then imported to Cytoscape (version 3.9.1; National Institute of General Medical Sciences, Bethesda, MD, USA) to visualize and classify protein networks of high cohesiveness. Statistical comparisons were calculated with two-way ANOVA or Students T-test with Holm-Sidak method using GraphPad Prism (v. 9.5.1). Level of significance was set at p < 0.05.

### 2.7 Mathematical modeling

We performed mathematical modeling using experimentally reported enzyme kinetics and kinetic parameters. The model and some of the parameters were based on previous models of dopamine homeostasis (Best et al., 2010, 2009) with some modifications to enable modeling of amino acid variation and to address reported data on dopamine storage and release in PC12 cells. The rate equations were implemented in Copasi (v4.35, (Hoops et al., 2006)) where the model simulations were run. Details about the model construction is provided in a supplemental section.

## 3 | Results

### 3.1 | High tyrosine and phenylalanine decrease dopamine levels in PC12 cells

To investigate the effects of phenylalanine and tyrosine levels on dopamine production, we exposed PC12 cells to different concentrations of these amino acids for periods of 1 to 6 and 1 to 24 hours, respectively. Serum phenylalanine levels 250-1000 µM are found in mild PKU (Van Wegberg et al., 2017), while the high concentrations of tyrosine used here (275-835 µM) reflect typical serum levels found in children treated with NTBC (Barone et al., 2020) and serum and brain tissue levels of NTBC treated mice (van Ginkel et al., 2022).

As shown in Fig. 1, high levels of phenylalanine rapidly depleted PC12 cells of tyrosine (Fig. 1A) and dopamine (Fig. 1B). This reduction was clearly observed already after 1 h, where the intracellular levels of tyrosine and dopamine were reduced by 77.2% (Fig. 1A) and 47.1% (Fig. 1B), respectively, in the presence of 1000 µM phenylalanine. After 6 h, the dopamine content was reduced by 20.3-69.5% in the presence of 100- 1000 µM L-Phe. This is in accordance with previous findings in cells and intact animals, showing that high concentrations of phenylalanine compete with the transport of other large neutral amino acids and can deplete the cells of neurotransmitter precursors (Pardridge, 1998). This also confirms that PC12 cells have an active exchange of metabolites with their extracellular environment and are dependent on a continuous supply of precursor amino acids to sustain their synthesis of catecholamines.

When cells were exposed to increasing concentrations of tyrosine, we observed a biphasic response: DA content was highest around physiological levels of tyrosine (75 µM), but gradually decreased from 3- 24 h at either at low (35 µM) or increasingly high (275 µM or 835 µM) concentrations of tyrosine (Fig. 1C). The viability and morphological features of the cells were unaffected even at the highest L-Tyr concentration (Fig. S2). Extracellular DA in the culture medium was also measured and although the concentrations of DA was much lower than in lysates, the data showed the same pattern, with DA levels being significantly decreased in the presence of both low and high concentrations of tyrosine (Fig. S3).

**Fig. 1.**
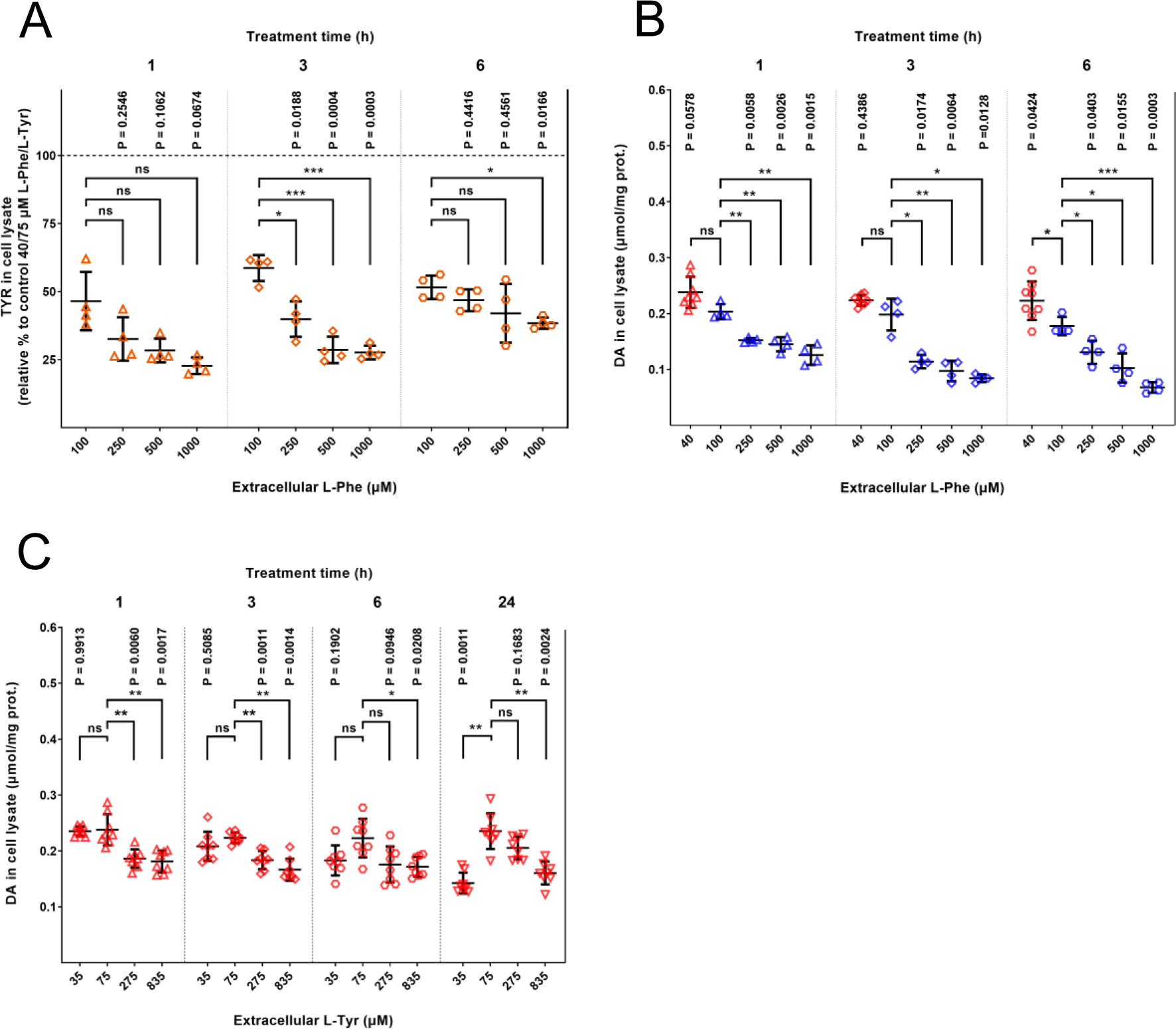
A) Tyrosine levels in PC12 cell lysates. Relative levels compared to cells exposed to 40 µM L-Phe and 75 µM L-Tyr (physiological), N=4; B) DA levels were measured in PC 12 cells exposed to different concentrations of extracellular L-Phe. All treatment conditions contained physiological (75 µM) L-Tyr in the media along with varying concentrations of L-Phe, N=4; C) Intracellular DA in cells exposed to cell culture media containing elevating L-Tyr levels. Physiological relevance is indicated for the varying treatment conditions on the X axis. Treatment conditions refer to extracellular concentration of tyrosine at the beginning of the treatment period and are as follows; Low L-Tyr: 35 µM; Physiological L-Tyr: 75 µM, moderately Increased L-Tyr: 275 µM and High L-Tyr: 835 µM. One-way ANOVA with Dunnett’s multiple comparisons tests were used to assess significance (P < 0.05). Error bars represent SD; N=8.

### 3.2 | Changes in extraneous L-tyrosine availability impacts TH phosphorylation

To explore how substrate availability could alter DA levels, we measured the concentration of TH protein and TH phosphorylation status in PC12 cells. Catecholamine synthesis is acutely regulated by levels of TH activation *via* phosphorylation on its N-terminal serines, particularly on the Ser19 and Ser40 residues (Ghorbani et al., 2020). PC12 cells were treated with media containing 35-835 µM L-tyrosine for either 1 or 6 hours. Once again, we observed a biphasic response to altered tyrosine concentrations. The total amount of TH protein was highest around 75 µM tyrosine but decreased at either lower (35-55 µM) or higher (115-835 µM) tyrosine concentrations (Fig. 2A). Intriguingly, these effects on TH protein concentrations were observed already at 1 h, indicating a rapid turnover of TH protein in these cells.

**Fig. 2.**
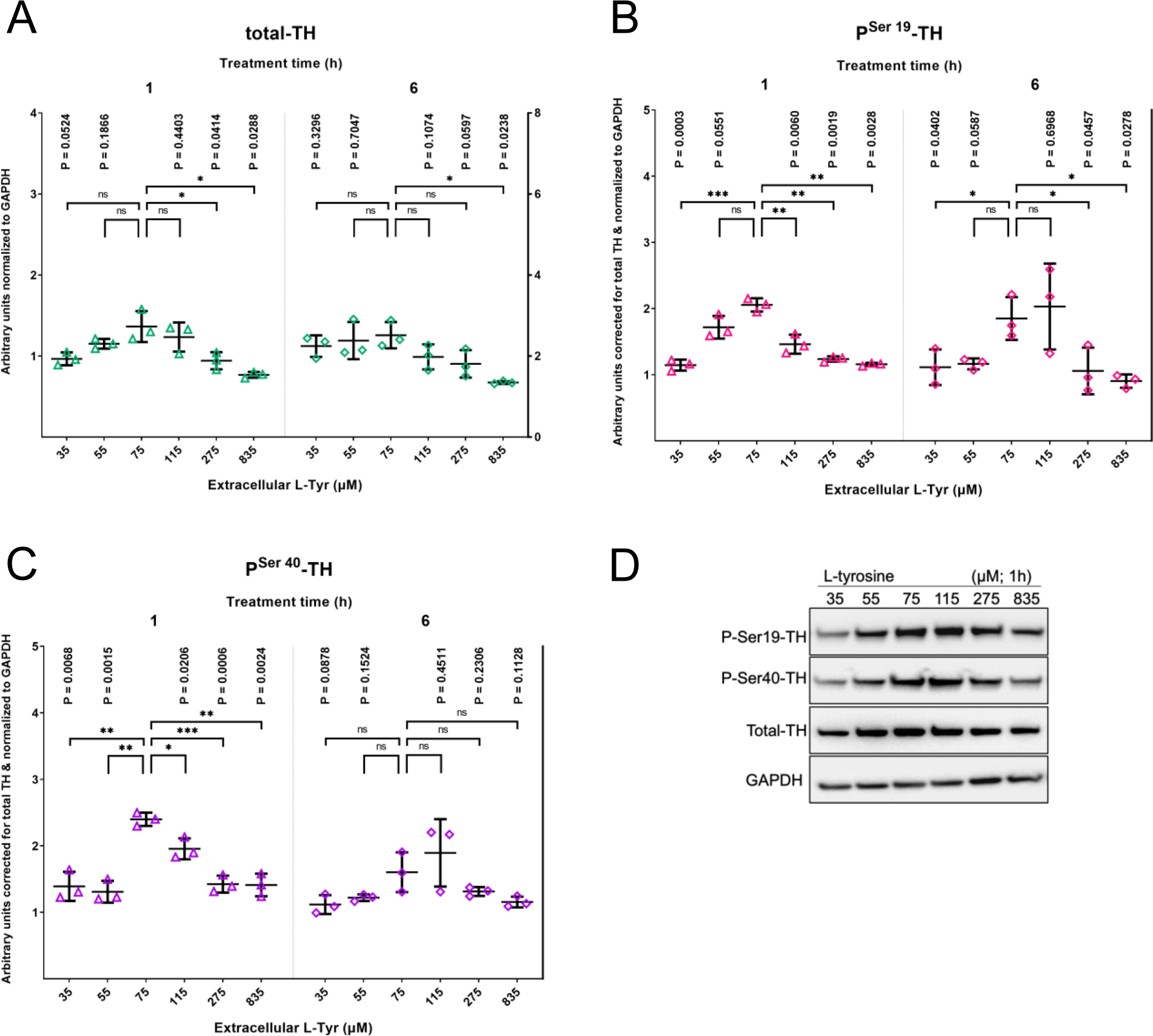
Effect of different tyrosine concentrations on TH protein levels and phosphorylation status. Immunoblot analysis of total TH protein (A) and TH phosphorylation at Ser19 (B), Ser40 (C) from PC12 cells exposed to different substrate concentrations (x axis) for 1 & 6 hours. Representative immunoblots (D). Student’s T-test with Welch’s correction was used to assess significance (two-tailed P-values; (P > 0.05)). Error bars represent SD; N=3.

Using antibodies targeting Ser19 and Ser40, we could observe that the site-specific phosphorylation at these sites (corrected for total TH protein) followed a similar pattern. For both sites, maximal TH phosphorylation (and by inference, probably also highest specific TH activity) was observed at or near physiological levels of tyrosine but was decreased under non-physiological conditions (Fig. 2B & C). This pattern of TH activation follows the alterations of dopamine content and might explain why dopamine levels are consistently lower outside the normal physiological range of tyrosine.

### 3.3 | Proteomic analysis identifies altered amino acid and monoamine metabolism in response to changing extracellular tyrosine concentration

To explore if tyrosine abundance could affect other cellular processes and protein levels, cells were incubated with either physiological (75µM), low (35µM), or high levels of tyrosine (275 and 835 µM) for 6 hours and subjected to a global proteomic analysis. Analysis of proteome changes showed that both low levels (35 µM) and very high levels of tyrosine (835 µM) were associated with many changes in protein profiles (Fig. 3A & B, respectively), whereas exposure to moderately elevated tyrosine (275 µM) was followed by only modest shifts (Fig. 3C).

**Fig. 3:**
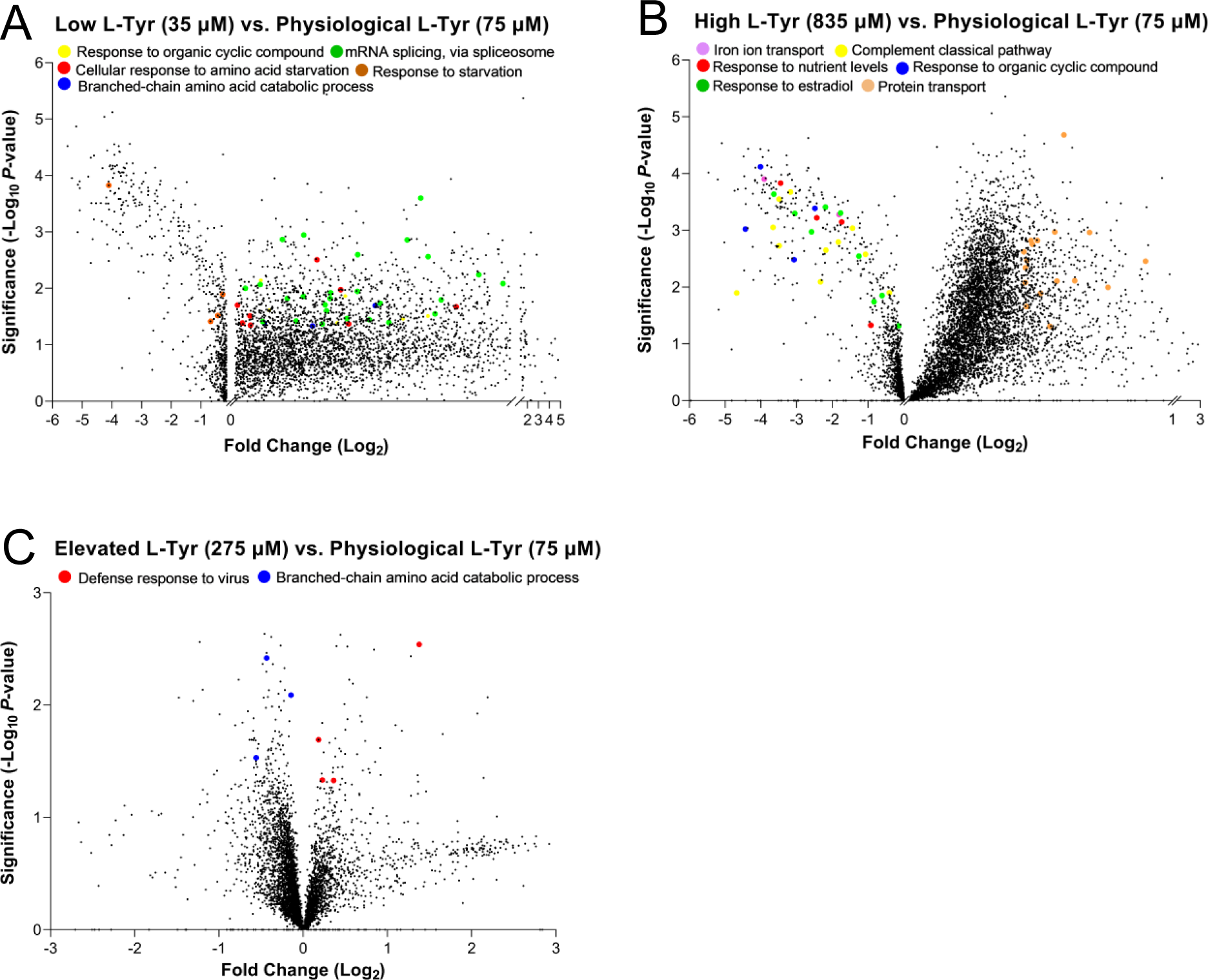
Volcano plots illustrate the shifting protein profiles responding to changes in substrate (L-Tyr) availability. Graphs A, B & C represent differentially abundant proteins plotted against P-values (-Log_10_) and Fold Change (Log_2_) that reach significance in comparisons between A) Low L-Tyr (35 µM) vs. physiological conditions, B) High L-Tyr (835 µM) vs. physiological conditions and C) moderately Elevated L-Tyr (275 µM) vs. physiological conditions. N=3; FDR < 0.05. Relevant proteins involved in the highlighted processes depicted on panels A & B are contained in Supplementary Table S1.

Altogether, when compared to physiological conditions, 1593 proteins were significantly differentially expressed in tyrosine-deprived cells (Fig 3A; 1257 upregulated and 336 downregulated), while 3286 proteins were significantly differentially expressed in cells supplemented with media containing high tyrosine levels (Fig 3B; 2972 upregulated and 314 downregulated). In comparison, few significant changes were observed at 275 µM tyrosine (Fig. 3C; 52 downregulated and 110 upregulated, i.e., a total of only 162 hits). For details, see Table S1, where proteins involved in multiple biological pathways represented in Fig. 3 are highlighted. For the complete list of significantly up- and downregulated proteins in response to environmental L-Tyr concentrations, please refer to Table S2 in a separate Excel spreadsheet.

Differentially expressed proteins were subjected to GO classification using the DAVID bioinformatics tool (<u>D</u>atabase for <u>A</u>nnotation, <u>V</u>isualization, and Integrated <u>D</u>iscovery). Proteins significantly enriched in PC 12 cells under either tyrosine depleted, or tyrosine supplemented conditions were primarily related to the catabolism of tyrosine and its byproducts. Key GO categories upregulated after tyrosine stimulation were amino acid catabolism, biosynthesis of cofactors, metabolism, protein-protein interaction, protein translation, and mRNA splicing.

Predominant biological processes that were found upregulated in response to low tyrosine levels included RNA binding, RNA splicing, branched amino acid catabolic process, cellular response to amino acid starvation and protein translation (Fig. 4A), suggesting that alternate anabolic mechanisms may have been initiated. GO analysis revealed processes that specifically involve TH activity, such as response to organic cyclic compounds, nutrient levels, and estradiol were downregulated at L-Tyr concentration of 835 µM (Fig. S4). This data suggests that high level of L-Tyr downregulates the above-mentioned pathways *via* reducing TH protein levels and activity.

**Fig. 4.**
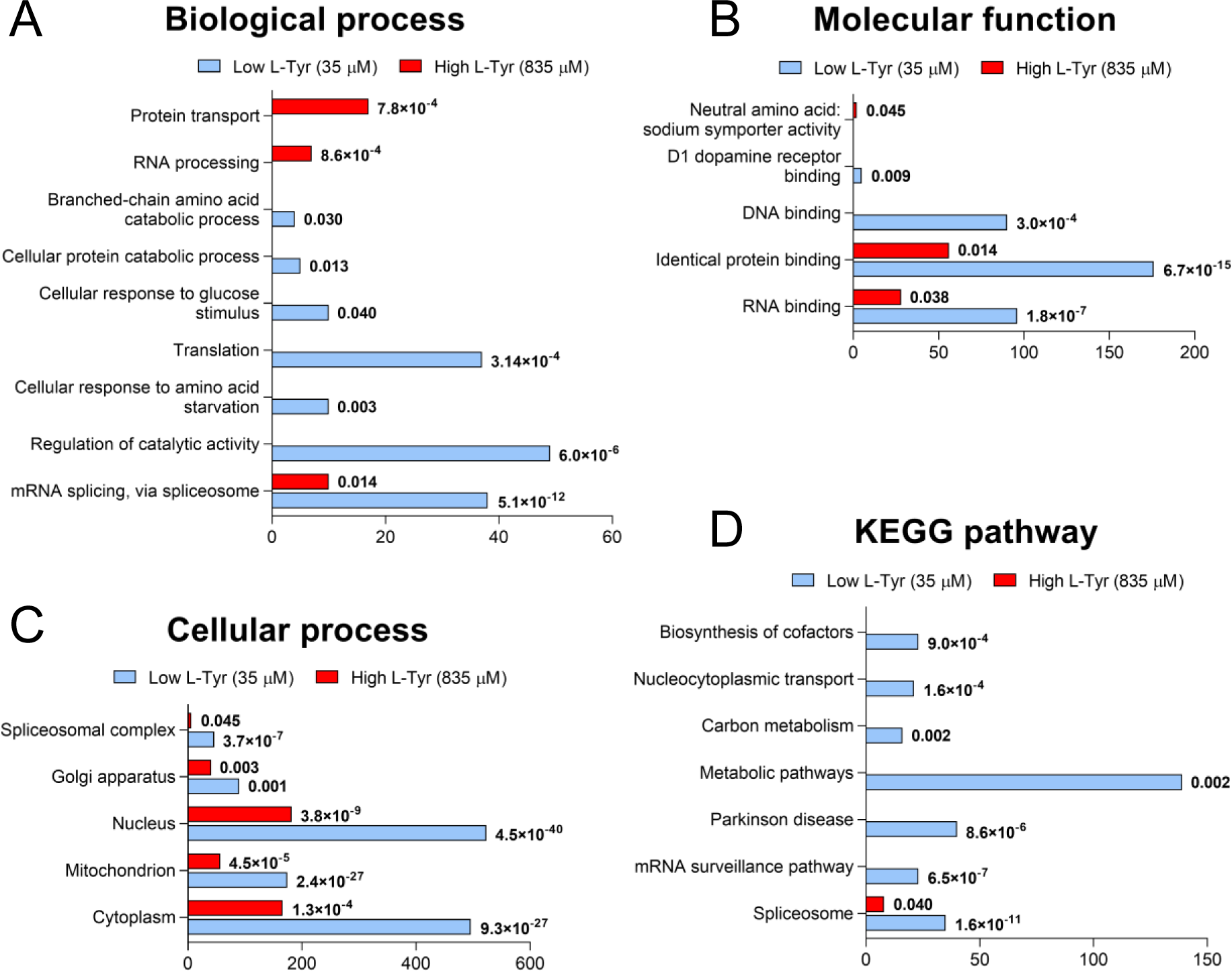
Unbiased proteomic analysis identified biological processes in PC-12 cells that responded to varying environmental L-Tyr. Gene ontology analysis of up-regulated genes under conditions mimicking hypotyrosinemia (35µM, blue) and high L-Tyr observed in treated TYRSN1 (835µM, red) are plotted by the following categories: Biological process (A), KEGG pathway analysis (B), Cellular component (C), and Molecular function (D). The X axes show the quantities of gene products with significantly altered abundance. P values are displayed next to each category (FDR < 0.05, 2-way ANOVA).

Our in-depth analyses have also identified several relevant proteins responding to non-physiological LNAA levels with altered expression levels. These include proteins involved in dopamine synthesis and metabolism (14-3-3α/β/λ, Dhfr, Qdpr, Spr, Pcbd1, Pcbd2, Comt, Maoa; Fig. 5A-B), amino acids transportation (Slc2a1/a4/a5, Slc7a5, Slc38a1/a2; Fig. 5E), branched amino acid catabolism (Bcat1, Bcat2, Ivd, Mcc1, Mccc2, Bckdha, Bckdhb; Fig. 5D), and dopamine receptor signaling (Gnai3, Gnao1, Gnaq; Fig. 5C). Importantly, several of these proteins are interaction partners of or are otherwise associated with TH at the center of DA biosynthesis (Fig. 5). Interestingly, proteins in the tyrosine catabolic pathway (Tat, Hppd, Hgo & Maai) were not detected to be significantly differentially expressed at high environmental L-Tyr levels. Intriguingly, however, the genes of both Lat1 (Slc7a5), the transmembrane transporter of L-Tyr, L-Phe and L-Trp as well as Bcat2, an enzyme in branched chain amino acid catabolism were significantly upregulated under that condition.

**Fig. 5:**
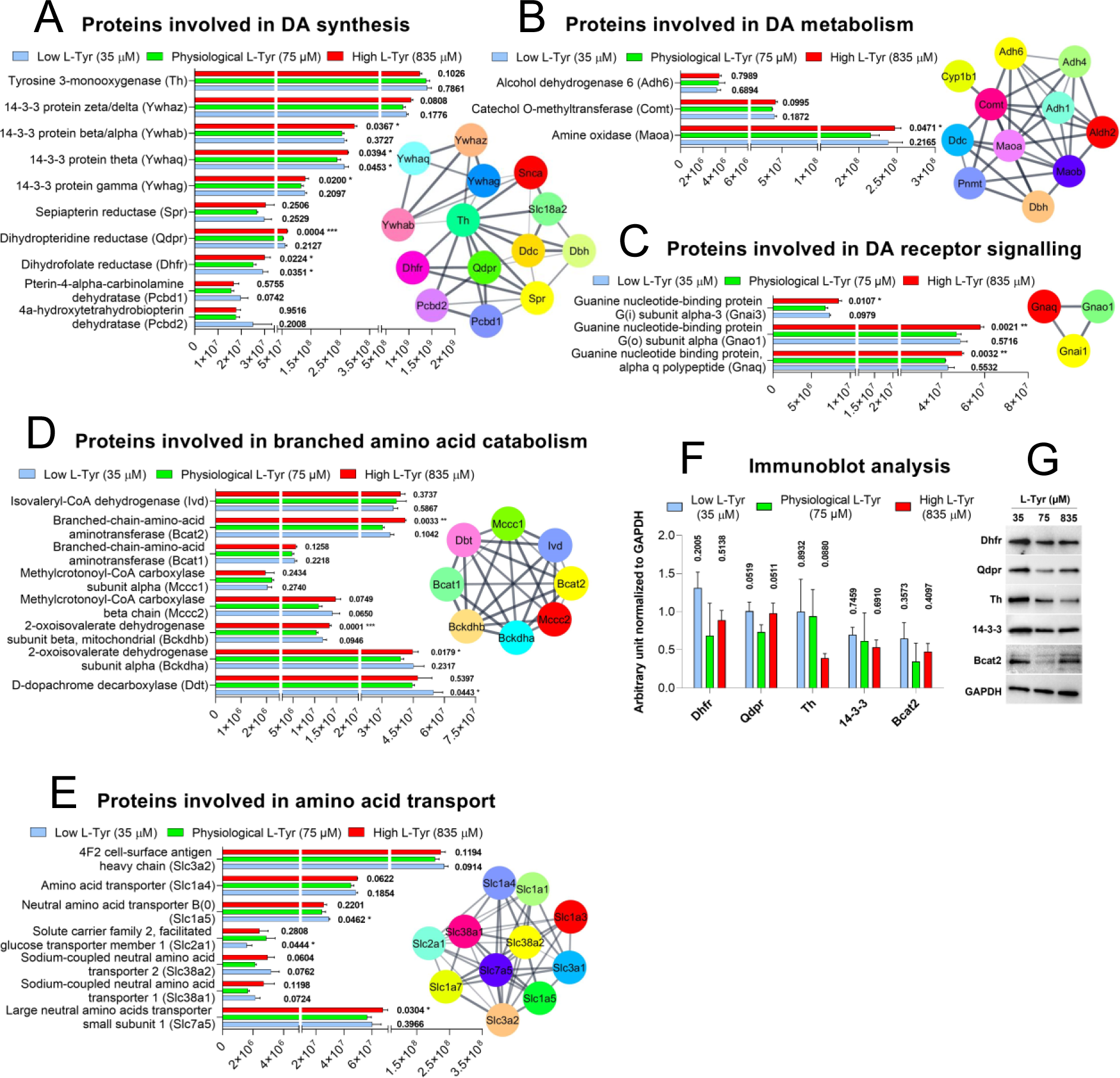
Deep proteomic analysis identified proteins with altered abundance in response to changing L-Tyr levels. PC12 cells maintained with low (35 µM, blue) or high L-Tyr (835 µM, red) containing media are compared to cells maintained under physiological condition (75 µM, green); plots show proteins involved in Dopamine synthesis (A), Dopamine metabolism (B), Dopamine receptor signalling (C), Branched amino acid catabolism (D), and Amino acid transport (E). Immunoblot analysis validates changes in the expression of Dhfr, Qdpr, 14-3- 3, Th and Bcat2 in lysates (F & G). Values are displayed as mean ± SD (2-way ANOVA, P < 0.05).

Relevant protein interaction networks have been generated using Cytoscape and were based on network topological information extracted from the STRING database (Aasebø et al., 2020). Proteins are depicted as nodes, while the strength of an interaction is signified by the thickness of connecting lines (Fig. 5A-E).

To complement the observed differences in protein expression levels obtained from deep proteomics- derived data, immunoblotting analysis against five key proteins, Dhfr, Qdpr, Th, 14-3-3, and Bcat2 were performed on PC12 lysates (Fig. 5F & G). We were able to verify the decrease of Th abundance at high L-Tyr levels and the effects of varying environmental L-Tyr on the other highlighted proteins as well.

### 3.4 Modelling of dopamine homeostasis under variable aromatic amino acid availability

We performed mathematical modelling to test the effects of different conditions on dopamine homeostasis in PC12 cells. The model development is described in detail in the supplemental text. We first modelled intracellular conditions where the cytosolic concentrations of Phe and Tyr were clamped (fixed) at different concentrations. This initial model used most of its parameter values from previous dopamine homeostasis models (Best et al., 2009) but had kinetic modifications to better address the impact of Phe and Tyr in PC12 cells. This model predicted a linear increase in time for vesicular refilling and decrease in cytosolic DA Conc. for increasing Phe concentrations (constant Tyr 75 μM) (Fig. S5C & D) and a bi-phasic relationship in response to Tyr conc. (constant Phe 100 μM) with highest cytosolic DA Conc. and fastest refilling rate at approx. 63 μM Tyr (Fig. S5E & F). However, this model did not predict any steady state change in vesicular DA and the model was far from capable of handling the reported spontaneous release of vesicular DA in PC12 cells or even the increase in vesicular pool size from cell growth. We therefore increased the catalytic capacity (V_max_-values) of the enzymes and transporters considerably to withstand an experimentally reported continuous spontaneous DA release rate of 0.7 % min^-1^ (Ritchie, 1979), but keeping the K_m_, K_i_ and K_si_ values the same.

The modified model has a higher steady state cytosolic DA (2.14 μM, Fig. 6A) prior to adding the continuous spontaneous DA release (Table S4), which then decreases to 16 nM (Fig. 6A). The Dopa steady state level increases from 175 nM prior to DA release to 1.74 μM with release included (Fig. 6A). As expected, the steady state level of vesicular DA decreased, but rather moderately from 79.9 to 72.4 mM after adding continuous release of vesicular DA (Fig. 6B) still well in line with experimental reports.

**Fig. 6.**
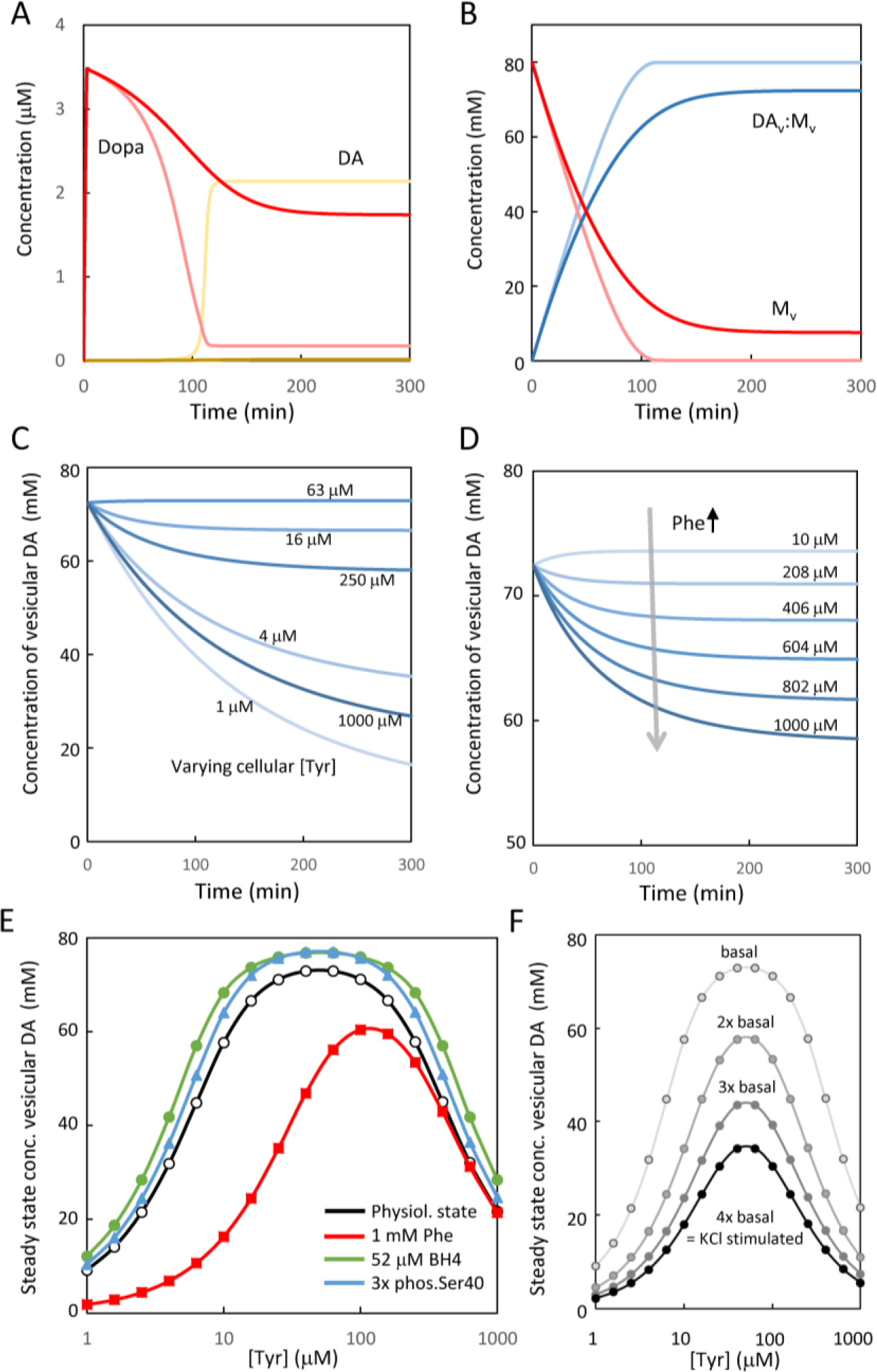
Mathematiccal modeling of dopamine homeostasis. In (A) and (B), the model was run from initial state (intracellular Phe and Tyr of 100 and 75 μM, respectively) without Dopa, cytosolic DA or vesicular DA until steady state was obtained. (A) shows the temporal change in Dopa (red) and cytosolic DA (yellow) in the absence (light colors) or presence (darker colors) of spontaneous vesicular DA release. (B) shows the temporal change in vesicular DA (DA_v_, blue) associated to the matrix (M_v_, red) for the model in the absence (light colors) or presence (darker colors) of spontaneous vesicular DA release. The majority of vesicular DA were in the form of matrix-bound DA (DA_v_:M_v_, 72.4 mM), whereas unbound vesicular DA remained low (0.956 μM). In (C) and (D), the situations were modelled where cells at steady state (Phe 100 μM, Tyr 75 μM) were challenged with a sudden change in the cellular Tyr (C) or Phe (D) concentration and the change in vesicular DA was followed over time (color coded from low (light) to high (dark) conc.). Panels (E) and (F) show the steady state conc. of vesicular DA at different cellular conc. of Tyr. In (E) the response to Tyr at basal conditions (black, Phe 100 μM, BH_4_ 26 μM, 5 % TH Ser40 phosphorylation) was compared to different cellular conditions where Phe (1 mM, red), BH_4_ (52 μM, green) and TH Ser40 phosphorylation (15 %, blue) was changed. In (F), the cellular conditions were in the normal state, but the rate of vesicular DA release was increased 2-4-fold that of basal spontaneous release rate and the steady state vesicular DA level was monitored as a function of cellular Tyr conc.

After including a continuous and spontaneous release of vesicular DA, the modelled steady state level of vesicular DA became sensitive to changes in TH activity responding to Tyr and Phe levels. Different situations were modelled where cells operating at physiological conditions were added media with altered Tyr (Fig. 6C) or Phe (Fig. 6D) levels. A linear decrease of vesicular DA was observed in response to increasing cellular Phe. A decrease in vesicular DA is predicted for Tyr levels of 1, 4, 16, 250 and 1000 μM, but a slight increase for 63 μM, suggesting that this concentration is slightly more optimal than 75 μM. Thus, a biphasic steady state level of vesicular DA is predicted in response to the cellular Tyr concentration. This relationship was further investigated in Fig. 6E, where the impact of increased BH_4_, Ser40 phosphorylation and high Phe (1 mM) were compared. The model predicted a steep increase in vesicular DA from 1-15 μM Tyr and a steep decrease from 160-1000 μM, whereas Tyr levels between 15-160 μM had only moderate impact on the vesicular DA levels. Doubling the cellular BH_4_ level increased the vesicular DA levels at all Tyr concentrations and broadened the tolerance range for Tyr. A similar but more moderate effect was found for increased Ser40 phosphorylation. The moderate effect of Ser40 phosphorylation is expected to be partly due to the low cytosolic DA levels in the model under continuous vesicular DA release. We further investigated the impact of increased continuous release rate of vesicular DA, 2 - 4x fold that of the basal spontaneous release (Fig. 6F), where a 4-fold increase corresponds to the reported release for KCl stimulated PC12 cells (Ritchie, 1979). Clearly, the increased release rate challenged DA homeostasis and the biosynthetic machinery even at optimal conditions for synthesis. Thus, at a 4-fold increased release rate relative to the basal, the model predicts a remaining vesicular DA level less than half that at basal conditions (under no reuptake).

We next extended the modeling to include transport of Tyr and Phe from the growth medium (see supplemental text and figures (Fig. S6 and S7) for details on implementation), using a similar, but much less extensive approach than Gauthier-Coles et al. (Gauthier-Coles et al., 2021) and experimental data from DePietro and Fernstrom (DePietro & Fernstrom, 1999). Our data were consistent with a major LAT1 (80 %) and minor LAT2 (20 %) mediated inward transport, but with an outward transport activity that was likely to involve other transporters (Fig. 7, S6, S7).

**Fig. 7.**
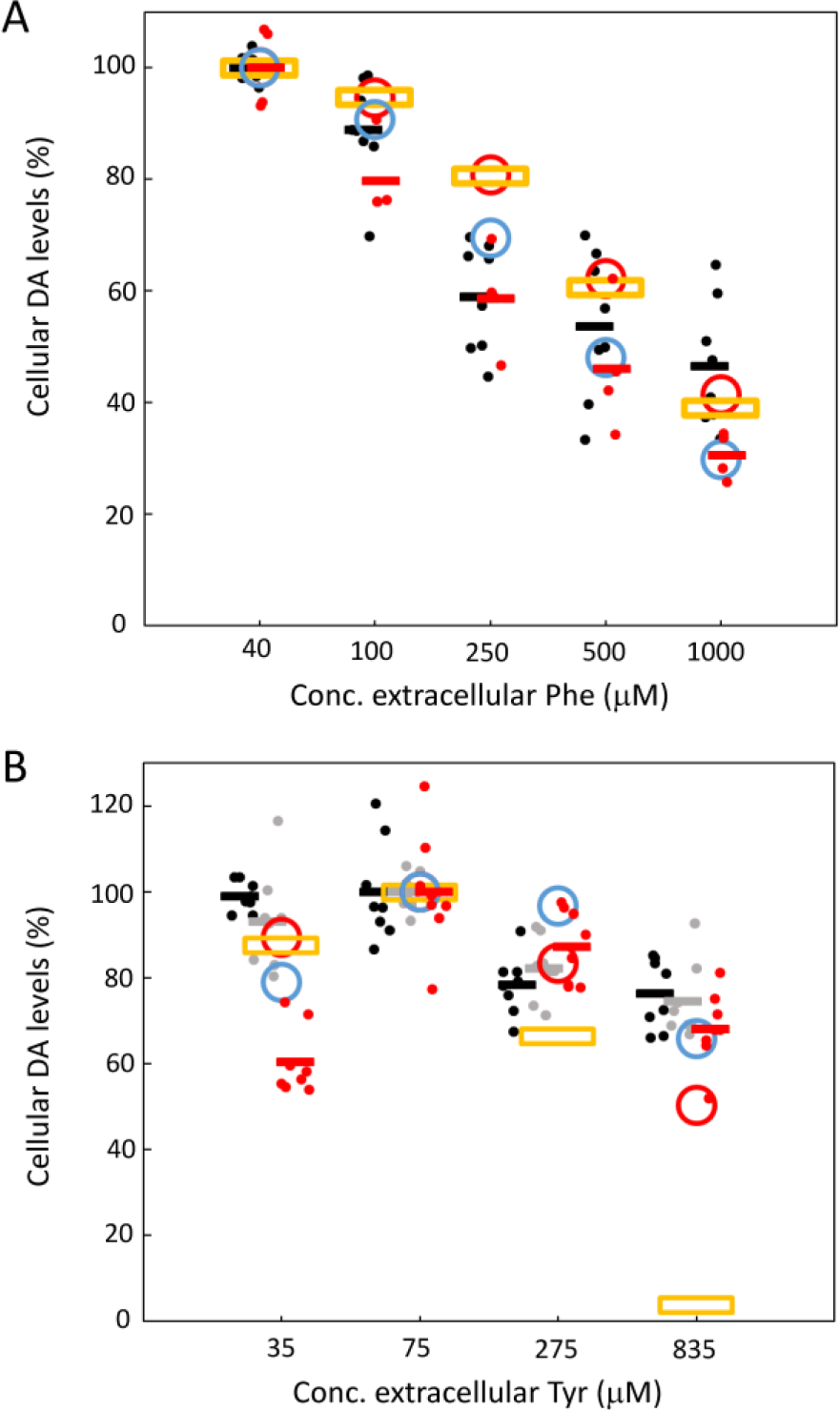
Comparison between experimental data and mathematical models. The cellular DA levels (shown as %) were compared between (A) data from Phe or (B) Tyr treatment of PC12 cells and that of model predictions for three models with different outward Tyr transport kinetics (low (orange); medium (blue); and high (red) V_max_ & K_m_, see supplemental text). In (A) data from 1 and 3 h treatments were pooled together (black dots, average as black line) and data from 6 h treatment is shown as red dots (average as red line). In (B) experimental data is shown as dots (black, 1 h; grey, 3 h; red, 24 h) with average values as lines.

It was clear that the cell growth conditions, and the underlying kinetics of cellular Tyr and Phe homeostasis would impact on how the DA synthesis pathway responded to changes in extracellular amino acid levels. We present the behavior of three amino acid transport situation where LAT1 and 2 dominate the inward transport and a different transporter with Michaelis Menten kinetics the outward transport. Three kinetic situations for outward transport (low, medium, and high V_max_ and K_m_, Fig. 7 and S7) were compared to experimental data from 1 and 3 h (Fig. 1A & B). Data from the Phe experiments fitted fairly well to all three models, but with the 6 h data being in best agreement with the medium V_max_, K_m_ model, which predicted the lowest Tyr_in_ levels (Fig. 7A). The higher agreement with the medium V_max_, K_m_ model was more evident in the Tyr experiments (Fig. 7B). We did not attempt a comprehensive fitting of the full model as our experimental data on amino acid transport were insufficient and this was beyond the scope of this paper. However, the experimental data and modeling together still point to TH kinetics being an important factor for understanding disturbances in monoamine homeostasis under conditions of dysregulated aromatic amino acid homeostasis.

## 4 | Discussion

Here we have shown that exposure of rat PC12 cells to either low or very high levels of tyrosine triggers rapid changes in the cellular proteome, including decreased protein levels and phosphorylation stoichiometry of TH, the rate limiting enzyme in DA synthesis. These changes were accompanied by an up to 39% decrease in levels of intra- and extracellular DA. Similarly, DA levels were decreased by up to 70% by increasing levels of extracellular phenylalanine. We used this data to develop a comprehensive model for how dopamine homeostasis is regulated by extracellular levels of aromatic amino acids. These findings may constitute as a novel pathophysiological response *in vivo*.

A motivation to perform these studies was to explore the molecular underpinnings of altered monoamine transmitter levels and brain function in the hereditary aminoacidopathies PKU and TYRSN1. For both conditions, a significant correlation has been reported between pathologically elevated amino acid levels and CNS symptoms. We recently reported such correlations between inattention symptoms, impaired working memory and plasma tyrosine levels (Barone et al., 2020). As NTBC treated TYRSN1 patients share cognitive symptoms with PKU and ADHD affected individuals, we proposed that a common pathophysiological mechanism is plausible. In this context, it is notable that central stimulants e.g. methylphenidate or amphetamines that increase prefrontal noradrenaline and DA transmission - can provide symptomatic relief in all three disorders (Barone et al., 2020). Hence, we argued that in humans, abnormally elevated tyrosine may limit catecholamine signaling in the PFC, partially *via* substrate inhibition of TH. By a similar mechanism, extremely elevated tyrosine and phenylalanine levels would probably affect serotonin biosynthesis as well through competitive inhibition of tryptophan transport and tryptophan hydroxylase activity (Barone et al., 2020).

Several hypotheses have aimed to explain the frequently occurring cognitive symptoms in NTBC treated TYRSN1, as well as tyrosinemia-2 and tyrosinemia-3, where high plasma concentrations of tyrosine are a common feature. The proposed mechanisms include a direct toxic effect of treatment with NTBC (a known herbicide), residual developmental brain damage from liver disease prior to treatment (Van Ginkel et al., 2016) and high plasma levels of tyrosine, either through direct toxicity *via* oxidative stress, as well as disrupted DNA-repair (De Prá et al., 2014; Macêdo et al., 2013) or indirectly due to the competitive nature of amino acid transport across the BBB where large neutral amino acids (LNAAs) use the same carrier (LAT1) (Pardridge, 1998). This has been confirmed in CSF samples of TYRSN1 patients, showing altered levels of several LNAAs (Thimm et al., 2011). Such an imbalance may alter catecholamine production, and may also decrease serotonin synthesis as well (Thimm et al., 2011). However, the suggested mechanisms on how elevated precursor levels affect DA synthesis and metabolism seem to diverge based on the experimental approaches utilized. Originally, the prevailing view was that a high supply of L-Tyr would result in elevated catecholamine synthesis and increased oxidative stress. This has been shown to occur under certain circumstances, specifically, in constantly firing dopaminergic retinal interneurons upon light exposure (Fernstrom & Fernstrom, 2007). However, as both too low and too high DA levels may produce cognitive symptoms, it has been difficult to establish whether elevated environmental tyrosine levels result in increased or attenuated DA synthesis (Barone et al., 2020; Thimm et al., 2011; van Ginkel et al., 2017).

Some studies concluded that artificially increased levels of L-Tyr raised L-Dopa production by TH without any apparent substrate inhibition (Brodnik et al., 2012; Wurtman et al., 1974), whereas others observed decreased catecholamine synthesis, possibly stemming from substrate inhibition (Badawy & Williams, 1982; Berger et al., 1996; DePietro & Fernstrom, 1998). To further complicate this picture, a recent study postulated that in the murine model of treated TYRSN1, monoamine neurotransmitter levels as well as behavioral and cognitive markers remained within normal parameters despite highly elevated tyrosine concentrations in the rat brain (van Ginkel et al., 2022). Earlier, Davison and colleagues arrived at a similar conclusion (Davison et al., 2019).

There can be many reasons for these diverging observations. At the protein level, kinetic properties of TH can differ between species, as can aromatic amino acid transporter levels and activation of relevant signaling pathways between cell types. Indeed, while tyrosine exerts substrate inhibition above 150 µM on rat Th (Fitzpatrick, 1991), human TH is inhibited at much lower tyrosine levels, falling within physiological concentrations (Blau et al., 1996) measured in plasma of healthy controls (Barone et al., 2020). Consequently, human patients may be more susceptible to pathologically elevated tyrosine levels characteristic to Tyrosinemias. Cellular BH_4_ homeostasis as well as free cytosolic DA levels will also greatly influence TH activity and the degree to which substrate inhibition can influence L-Dopa synthesis (see Fig. 6). Some researchers have studied L-Dopa synthesis in the presence of a Dopa Decarboxylase inhibitor (NSD-1015), whereas others have reported DA levels or DA metabolite levels. The comparison of results using these different readouts is not trivial (see below). Critically, the effective extracellular concentration of tyrosine surrounding brain dopaminergic nerve terminals remains largely unknown and may fluctuate depending on dietary protein intake and the concentration of other competing metabolites. Uptake of L-Tyr is shown to increase proportionally to elevated external concentrations. Some recent animal studies show a close correlation between plasma and tissue levels of tyrosine (Davison et al., 2019; van Ginkel et al., 2022), however, some earlier studies on animals indicated that brain tyrosine concentrations do not exactly match plasma levels of L-Tyr (Berger et al., 1996; Harding et al., 2014).

It has been proposed, based on mathematical modelling, that substrate inhibition may play a distinct role in monoamine transmitter homeostasis (Best et al., 2009; Reed et al., 2010) compensating for fluctuating precursor availability following varying dietary intake (Fernstrom & Fernstrom, 1994). Such regulatory mechanism may be advantageous for cellular viability by limiting direct and indirect metabolic costs to catecholamine production (Meiser et al., 2013). Our model confirms this robustness of vesicular DA levels for Tyr_in_ between 10-250 µM, which could be further strengthened by Ser40 phosphorylation or increasing BH_4_ levels. Phe is known to increase BH_4_ in the liver, but it is not known whether this also occurs in dopaminergic cells or if L-Tyr has a similar effect on BH_4_ levels. Challenging cells with increased continuous DA release seems to break down this robustness, which could suggest that DA deficiency becomes more prevailing in high activity states.

Our experimental findings also appeared to be in congruence with the hypothesis that in catecholaminergic cells, DA synthesis may be negatively affected at pathologically elevated L-Tyr levels. Unbiased proteomic analysis demonstrated a wide-ranging, nuanced cellular response with 336, 52 & 314 significantly down-regulated and 1257, 110 & 2972 up-regulated proteins identified, respectively, when L-Tyr-deprived, and L-Tyr-over-stimulated cells (from the lower to the higher range of the pathological spectrum) typical to treated TYRSN1 were compared to cells maintained under physiological conditions. Among the notable findings were interaction partners of TH; variants of 14- 3-3, proteins involved in its co-factor biosynthesis (Dihydropteridine reductase, Dhpr), DA metabolism (Catechol O-methyltransferase, Comt; Monoamine oxidase A, Maoa), DA-receptor signaling, amino acid transport as well as amino acid catabolic processes. Interestingly, proteins in the tyrosine catabolic pathway (Tat, Hppd, Hgo & Maai) were not detected as significantly altered under high environmental L-Tyr levels. On the other hand, it was notable that both the Lat1 (Slc7a5) and Bcat2 genes were significantly upregulated under that condition, indicating possible compensatory responses. Lat1 is the cross-membrane transporter of branched-chain and bulky amino acids (including phenylalanine and tyrosine), while the latter is a critical enzyme in branched chain amino acid catabolism. TH expression itself was significantly reduced at highly elevated substrate levels, corroborated also by immunoblotting experiments that indicated corresponding site-specific changes in TH activation to occur *via* decreased phosphorylation on both Ser19 & Ser40 regulatory sites, however, these changes were not significant for S40 after 6 hours. In this context, it is notable that modest L-Tyr stimulation (0.5 µM) has been shown to induce Ser40 phosphorylation, resulting in enhanced TH activity in kidney cells (Taveira-Da- Silva et al., 2019), however, to our knowledge, similar experiments with extreme L-Tyr levels have not been performed prior to our efforts.

We detected rapid and significant declines of intracellular DA content under non-physiological conditions. Extracellular DA from the media after treatment was also measured and although detection of DA was low, we registered the same biphasic response (Fig. S3). Results showing how at the earlier time points, only high substrate levels were shown to significantly reduce DA levels (Fig. 1C) - despite similar changes seen in the turnover and activation of TH to those observed under hypotyrosinemic conditions (Fig. 2) - seem to reaffirm that; a) TH operates at (or above) saturation levels of its substrate under physiological conditions (Kaufman, 1995), thus being able to maintain quasi normal DA synthesis for a short time during substrate deprivation and b) that substrate inhibition may play a direct role *in vivo* in preventing runaway DA synthesis in the short term. Over time, while DA production in PC12 treated with 275 µM L-Tyr appeared to slowly normalize, the decrease remained significant at 835 µM and reached significance under hypotyrosinemic conditions after 24 hours. This suggests that if plasma levels of L-Tyr can be maintained in treated HT-1 patients under 300 µM or preferably lower, potential precursor-related disturbances in DA levels may be partially alleviated. Indeed, some ADHD related symptoms were less prominent in patients with plasma L-Tyr levels under 400 µM (Barone et al., 2020).

Phenylalanine is a substrate of TH in chromaffin cells, PC12 cells and intact animals (Fukami et al., 1990) and competes with tyrosine for amino acid transport and metabolism, including catecholamine synthesis. Elevating phenylalanine levels appear to disrupt DA production *via* competition at the plasma membrane LNAA transporter (Fig. 1B). As tryptophan uses the same transporter, the same mechanism may be responsible for compromising serotonin biosynthesis (Lykkelund et al., 1988). Notably, tyrosine uptake was found to be much more sensitive to changes in the extracellular balance within the LNAA pool (Fig. 1A) than DA production, further signifying the regulatory importance that TH operates above saturating precursor concentrations. Consequently, aside from keeping plasma concentration of L-Tyr under 300 µM, the goal should be to maintain a physiologically relevant LNAA ratio to prevent deficiencies in other pathways involving these amino acids. The modeling shows a close relationship between cellular growth conditions, the amino acid transport kinetics, and the response to amino acid disturbances in the dopamine synthesis. Understanding amino acid homeostasis in monoamine producing cells is therefore likely to support our understanding of neuropsychiatric symptoms in metabolic disorders with highly dysregulated circulatory amino acid levels.

### 4.3 | Methodological considerations

The relevant *in situ* cellular concentrations and dynamics of amino acid transport and catecholamine synthesis are only partially understood. While *in vitro* experiments consistently show that substrate inhibition is an inherent property of TH (Barone et al., 2020; Quinsey et al., 1998; Szigetvari et al., 2019; Tekin et al., 2014), conclusions drawn from *in vivo* studies using cell cultures and live animals have been inconsistent. The latter approach has often relied on using high concentrations of the DOPA-decarboxylase inhibitor NSD-1015 (Brodnik et al., 2012; DePietro & Fernstrom, 1999; Westerink et al., 1990). However, NSD-1015 is a non-specific and incomplete inhibitor of DOPA-decarboxylase (Bongiovanni et al., 2006), and although these studies were better able to discern direct changes in L-Tyr hydroxylation, the limitations of introducing artificial blockers in such a tightly regulated metabolic pathway may also lead to artefacts. We also find that the elimination of DA feedback inhibition and other downstream feedback mechanisms that control TH activity may constitute a major limitation in studying catecholamine metabolism.

A strength of our study is that we aimed to mimic a natural cellular environment by using complex, serum-rich media also during treatment and by avoiding specifically targeting certain enzymes within the chain of metabolic activity. As a metabolic intermediate, L-DOPA is highly transient in nature and at any selected time, its concentration was too low for accurate measurements (Fig. S1). Instead, we measured total intracellular dopamine content. In dopaminergic cells, the vast majority (∼97%) of synthetized DA is sequestered in vesicles (Li et al., 2018) and this vesicular content would be emptied in an hour under “normal” firing rate were it not for continued synthesis and a parallel DA re-uptake process orchestrated by DAT (Kadota et al., 1996). According to our data, extracellular DA gradients followed similar course to those measured intracellularly - that is, reduced DA content at too low (35 µM) or too high (275 & 835 µM) L-Tyr levels (Fig. S3). Therefore, we can infer that most of the decrease originates from disrupted synthesis and to a lesser extent, catabolic processes.

PC12 cells are derived from rat chromaffin cells (pheochromocytes). As chromaffin cells constitute the largest and most homogenous collection of catecholamine synthesizing cells in mammalian tissues, primary chromaffin cells and PC12 cells are preferred model systems for mechanistic studies of DA synthesis-related cellular processes, including regulation of TH activity (de Siqueira et al., 2023; Lisek et al., 2022). Our studies are comparable to previous investigations on the effects on elevating L-Phe and L-Tyr levels in PC12 cells (DePietro & Fernstrom, 1998). However, not all findings in cultured cells are directly transferable to intact tissues or organisms. Thus, it would be interesting to compare our findings to studies on different populations of catecholaminergic cells derived from the human brain. However, the brain contains hundreds to thousands of different cell types, each with their unique transcriptomic profiles and specific spatial and biochemical interactions with neighboring cells (Tasic et al., 2019). Thus, even for brain derived tissues, cells or cell lines, it is challenging to extrapolate from a model system to intact tissues or whole organism.

## 5 | Conclusion

Using PC12 cells as model system, we have demonstrated that these cells respond to increased tyrosine substrate availability by a rapid downregulation of their DA production. This paradoxical effect appears to be mediated by multiple mechanisms, including substrate inhibition of TH, downregulation of TH protein and phosphorylation levels, altered regulation of amino acid transporters (including Lat1), as well as widespread changes in intracellular amino acid and monoamine metabolism. The experimental data and mathematical modeling point to TH kinetics as being particularly important for understanding disturbances in monoamine homeostasis under conditions of dysregulated aromatic amino acid homeostasis, such as in PKU and TYRSN1. From a clinical perspective, it may be necessary to avoid extremely high levels of plasma tyrosine and to use tryptophan supplementation in TYRSN1 patients treated with NTBC (Barone et al., 2020; Thimm et al., 2011). New technologies and approaches may be needed to fully understand the consequences of altered amino acid levels on brain metabolism and functions, thus improving our understanding of the underlying pathophysiological processes and management of patients with aminoacidopathies and other (neuro)metabolic disorders.

## ACKNOWLEDGEMENTS

This work was supported by grants from Stiftelsen Kristian Gerhard Jebsen (SKGJ-MED-02), The Regional Health Authority of Western Norway (912264), The Research Council of Norway the European Union’s Horizon 2020 research and innovation programme (CoCA), The Norwegian ADHD Research Network for funding.

## AUTHOR’S CONTRIBUTIONS

Peter D. Szigetvari and Jan Haavik contributed to study design. Peter D. Szigetvari wrote the original draft, developed methods for cell culture studies and performed various *in vitro* experiments, statistical analyses as well as data interpretation. Sudarshan Patil also performed various *in vitro* experiments and together with Even Birkeland, they provided proteomic assessment. Rune Kleppe contributed to writing the article and provided extensive mathematical modelling. Jan Haavik supervised and coordinated the study, also contributed to writing the manuscript as well as provided critical review before submission. All authors read, reviewed, and approved the final manuscript.

## DATA AVAILABILITY STATEMENT

The data that supports the findings laid out in this work are available from the corresponding author upon reasonable request. Statistical reports for all datasets are included in a separate Excel spreadsheet.

