## Supplementary material for "The effects of phenylalanine and tyrosine levels on dopamine production in rat PC12 cells. Implications for treatment of phenylketonuria and tyrosinemia type 1"

*Jan Haavik* 0000-0001-7865-2808

*Peter D. Szigetvari* 0000-0002-1821-2779

*Sudarshan Patil* 0000-0002-6294-7959

*Rune Kleppe* 0000-0002-6086-755X

*Even Birkeland* 0000-0002-4605-0762

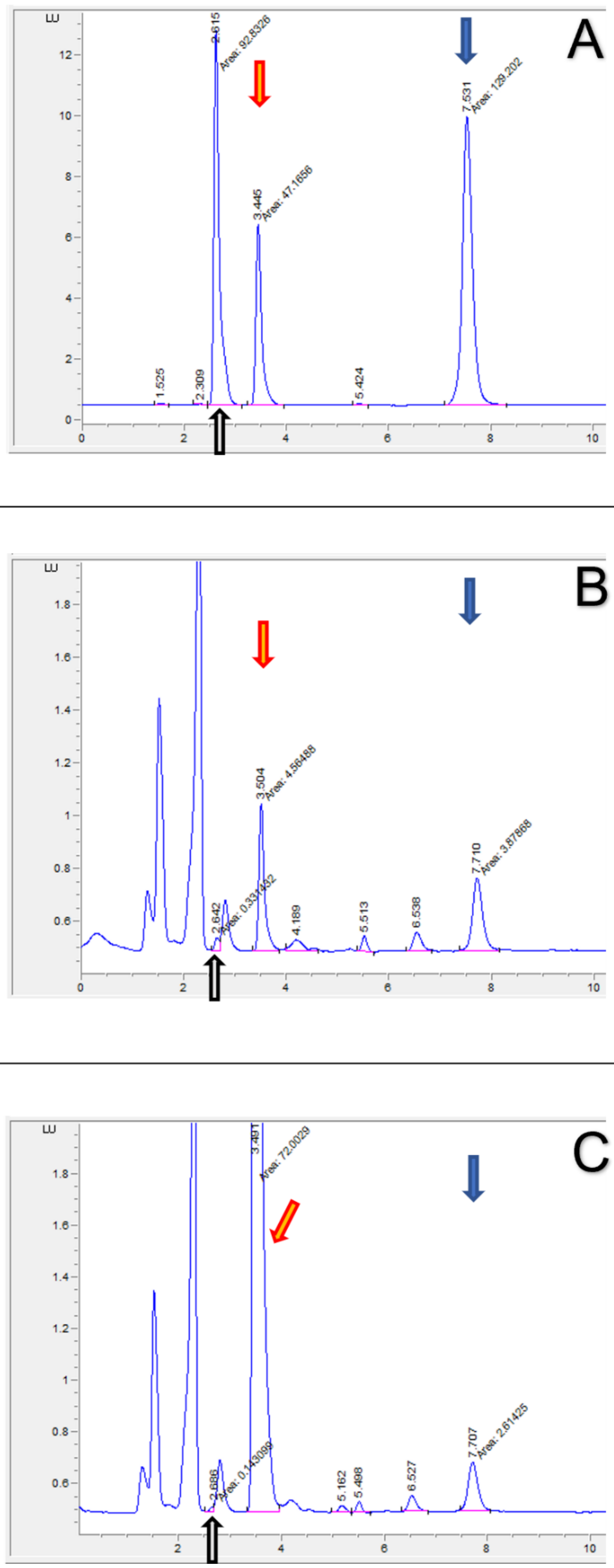


Fig. S1. Representative chromatograms displaying peak separations of L-Dopa, L-Tyr and DA both in samples containing pure standards (A); and lysates (B) Treated PC12 (1h-75 µM L-Tyr) & (C) Treated PC12 (1h-835 µM L-Tyr). Color coding: grey: L-Dopa, red: L-Tyr, blue: DA.

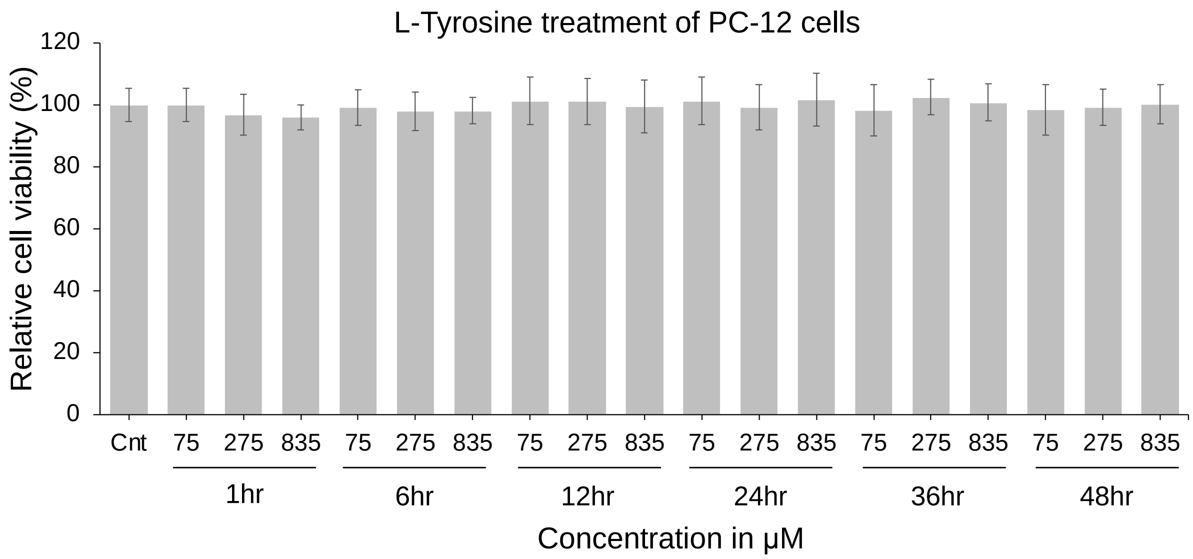


*Fig. S2. Cellular viability over the course of 48 hours under experimental L-Tyr treatment conditions were assessed. We found no significant effect on cellular survivability in response to either varying L-Tyr concentrations or elapsed time (within acceptable margins for cultured PC12 cells, that is).*

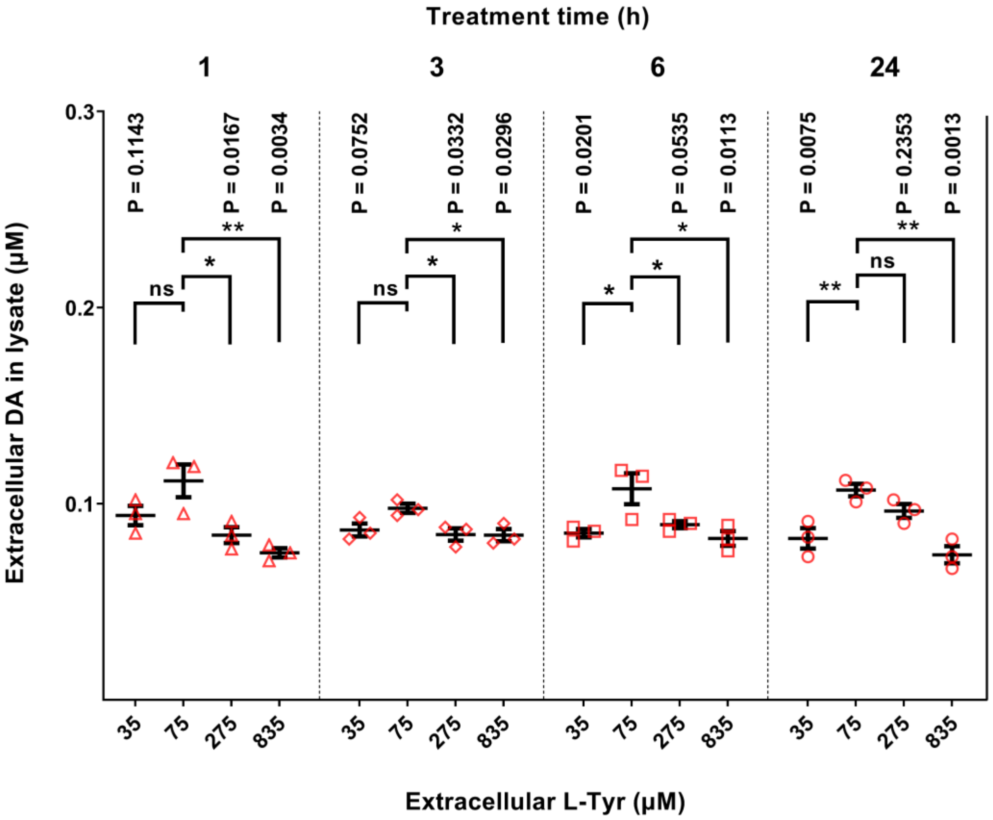


*Fig. S3. PC12 cells exposed to cell culture medium containing elevating concentrations of L-Tyr. Physiological relevance is indicated for the varying treatment conditions on the X axis. Treatment conditions refer to extracellular concentration of tyrosine at the beginning of the treatment period and are as follows Low tyrosine: 35 µM; Physiological: 75 µM, moderately Increased tyrosine: 275 µM and High tyrosine: 835 µM. 2-way ANOVA was used to assess significance. Error bars represent SD; N=3 (technical replicates).*

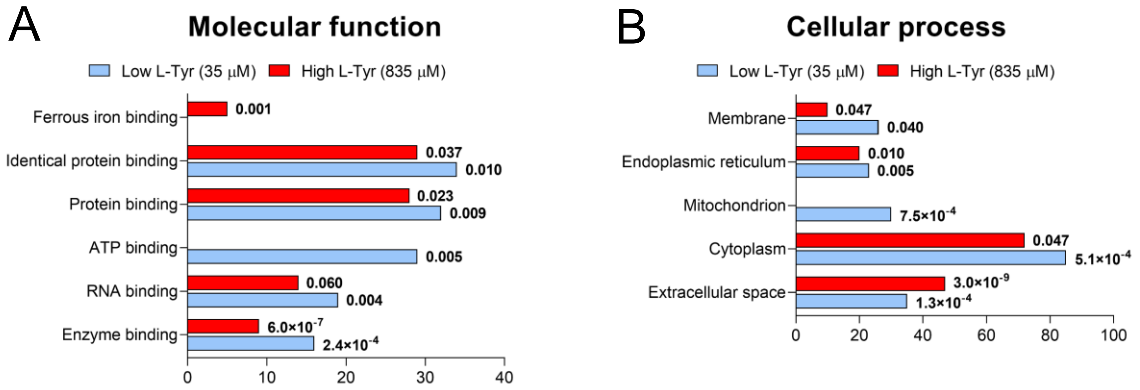


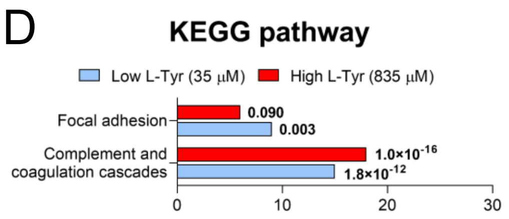


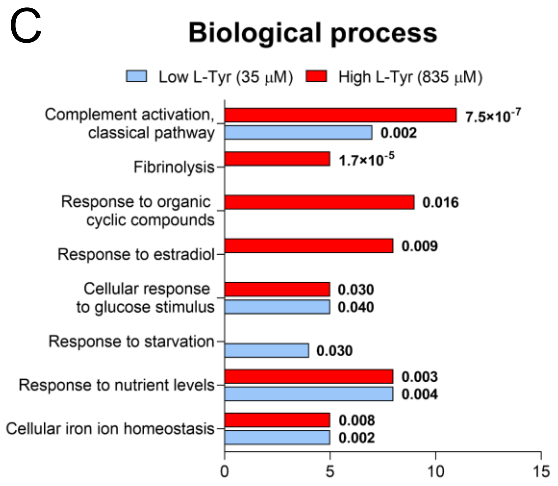


*Fig. S4. Proteomic analysis identified biological processes responding to varying environmental L-Tyr levels. Gene ontology analysis of downregulated genes in L-Tyr deprived- (35μM; blue) and L-Tyr overstimulated cells (835μM; red); plots for Molecular function (A), Cellular process (B), Biological process (C), and KEGG pathway analysis (D) with number of genes in each category ($p < 0.05$, 2-way ANOVA).*

| Low L-Tyr (35μM) Vs Physiological (75μM) | | | |
|---|---|---|---|
| **Accession no** | **Fold change (Log2)** | **P-Value(<0.05)** | **Protein description** |
| | **Response to Starvation** | | |
| P06866 | -4.1087 | 0.0001490 | Haptoglobin (Hp) |
| F1LM16 | -0.6752 | 0.0383398 | Plasminogen activator inhibitor 1 (Serpine1) |
| D3ZMG0 | -0.4382 | 0.0302985 | Serine/threonine-protein kinase (Ulk1) |
| Q6P2A5 | -0.2655 | 0.0127684 | GTP:AMP phosphotransferase AK3, mitochondrial (Ak3) |
| **Cellular response to organic substance** | | | |
| P63086 | 0.1784 | 0.0423464 | Mitogen-activated protein kinase 1 (Mapk1) |
| P55213 | 0.2007 | 0.0071752 | Caspase-3 (Casp3) |
| P18418 | 0.2105 | 0.0244915 | Calreticulin (Calr) |
| Q6AYN8 | 0.2920 | 0.0409087 | DNA replication licensing factor MCM7 (Mcm7) |
| F1M9D6 | 0.3016 | 0.0136663 | Signal transducer and activator of transcription (Stat1) |
| P30121 | 0.3708 | 0.0345731 | Metalloproteinase inhibitor 2 (Timp2) |
| D3ZCS4 | 0.4005 | 0.0308057 | IQ motif-containing GTPase-activating protein 3 (Iqgap3) |
| P07825 | 0.5429 | 0.0269934 | Synaptophysin (Syp) |
| **Cellular response to amino acid starvation** | | | |
| A0A0H2UHU0 | 0.1727 | 0.0198125 | 40S ribosomal protein S25 (Rps25) |
| A0A0G2K714 | 0.1786 | 0.0411876 | Endophilin-B1 (Sh3glb1) |
| P68101 | 0.1875 | 0.0305932 | Eukaryotic translation initiation factor 2 subunit 1 (Eif2s1) |
| Q63184 | 0.1882 | 0.0449922 | Interferon-induced, double-stranded RNA-activated protein kinase (Eif2ak2) |
| D3ZZH6 | 0.2677 | 0.0031002 | Mitogen-activated protein kinase kinase kinase (Map3k5) |
| Q91XJ1 | 0.2960 | 0.0105728 | Beclin-1 (Becn1) |
| Q63486 | 0.3056 | 0.0425870 | Ras-related GTP-binding protein A (Rraga) |
| B2GUV6 | 0.4344 | 0.0211479 | RCG31441 (Sesn2) |
| Q9JHE5 | 0.6924 | 0.0207444 | Sodium-coupled neutral amino acid transporter 2 (Slc38a2) |
| **Branched-chain amino acid catabolic process** | | | |
| O35854 | 0.1687 | 0.0294566 | Branched-chain-amino-acid aminotransferase, mitochondrial (Bcat2) |
| A0A0A0MXW1 | 0.2049 | 0.0412596 | 2-oxoisovalerate dehydrogenase subunit beta, mitochondrial (Bckdhb) |
| Q5XIE6 | 0.2623 | 0.0464344 | 3-hydroxyisobutyryl-CoA hydrolase, mitochondrial (Hibch) |
| Q6P6R2 | 0.3374 | 0.0201391 | Dihydrolipoyl dehydrogenase, mitochondrial (Dld) |
| **mRNA splicing, via spliceosome** | | | |
| D4A997 | 0.1664 | 0.0124858 | HIV TAT specific factor 1 (Predicted) (Htatsf1) |
| D3ZAS8 | 0.1820 | 0.0099939 | Spliceosome-associated factor 3, U4/U6 recycling protein (Sart3) |
| M0R835 | 0.2000 | 0.0084968 | Splicing factor 3B, subunit 6 (Sf3b6) |
| G3V9M1 | 0.2028 | 0.0387579 | RNA helicase (Ddx23) |
| B5DES0 | 0.2265 | 0.0013532 | Small nuclear ribonucleoprotein Sm D2 (Snrpd2) |
| D4A2C6 | 0.2323 | 0.0150756 | U6 snRNA-associated Sm-like protein LSm4 (Lsm4) |
| A0A0G2K3S6 | 0.2430 | 0.0379073 | RNA binding motif protein 10, isoform CRA_a (Rbm10) |
| E9PT66 | 0.2516 | 0.0137876 | Splicing factor 3b, subunit 3 (Sf3b3) |
| F1LM66 | 0.2521 | 0.0011288 | 116 kDa U5 small nuclear ribonucleoprotein component (Eftud2) |
| F1LSW6 | 0.2738 | 0.0427800 | Inositol hexakisphosphate and diphosphoinositol-pentakisphosphate kinase (Ppip5k1) |
| F1LM37 | 0.2775 | 0.0195048 | Splicing factor 1 (Sf1) |
| Q9JMJ4 | 0.2790 | 0.0248438 | Pre-mRNA-processing factor 19 (Prpf19) |
| F1LNJ2 | 0.2834 | 0.0150876 | U5 small nuclear ribonucleoprotein 200 kDa helicase (Snrnp200) |
| D3ZZR5 | 0.2840 | 0.0119909 | Small nuclear ribonucleoprotein polypeptide A' (Snrpa1) |
| P55770 | 0.3035 | 0.0340380 | NHP2-like protein 1 (Snu13) |
| A0A0G2JTI7 | 0.3160 | 0.0112695 | Pre-mRNA-splicing factor 3 (Prpf3) |
| G3V7T6 | 0.3164 | 0.0025248 | Splicing factor 3b, subunit 1 (Sf3b1) |
| Q5FVI5 | 0.3305 | 0.0358301 | Thioredoxin-like protein (Txnl4b) |
| Q63413 | 0.3433 | 0.0183065 | Spliceosome RNA helicase Ddx39b (Ddx39b) |
| O70454 | 0.3538 | 0.0402322 | Protein BUD31 homolog (Bud31) |
| A0A0G2JXW4 | 0.3756 | 0.0013781 | Heterogeneous nuclear ribonucleoprotein C, isoform CRA_a (Hnrnpc) |
| Q6AYL5 | 0.3862 | 0.0039207 | Splicing factor 3B subunit 4 (Sf3b4) |
| D3ZMS1 | 0.3918 | 0.0002504 | Splicing factor 3b, subunit 2 (Sf3b2) |
| D3ZQM0 | 0.4008 | 0.0027441 | Splicing factor 3a, subunit 1 (Sf3a1) |
| D4A7J8 | 0.4091 | 0.0282975 | PRP4 pre-mRNA processing factor 4 homolog (Yeast) (Prpf4) |
| Q4KLH4 | 0.4163 | 0.0159133 | Paraspeckle component 1 (Pspc1) |
| Q6AYK1 | 0.4614 | 0.0056897 | RNA-binding protein with serine-rich domain 1 (Rnps1) |
| Q27W01 | 0.4903 | 0.0081990 | RNA-binding protein 8A (Rbm8a) |
| Q4KLI7 | 0.5616 | 0.0221423 | Splicing factor 3a, subunit 3 (Sf3a3) |
| O08837 | 0.5786 | 0.0062369 | Cell division cycle 5-like protein (Cdc5l) |
| Q6AXT8 | 0.5788 | 0.0193575 | Splicing factor 3A subunit 2 (Sf3a2) |
| P83871 | 0.6499 | 0.0452387 | PHD finger-like domain-containing protein 5A (Phf5a) |
| D4A5T1 | 0.6870 | 0.0455004 | Splicing factor 3B subunit 5 (Sf3b5) |
| Q99M63 | 0.6882 | 0.0007779 | WD40 repeat-containing protein SMU1 (Smu1) |
| B2GV05 | 0.7906 | 0.0314452 | RNA-binding protein 5 (Rbm5) |
| D4ACM9 | 0.9652 | 0.0000120 | Microfibrillar-associated protein 1A (Mfap1a) |

| High L-Tyr (835μM) Vs Physiological (75μM) | | | |
|---|---|---|---|
| **Accession no.** | **Fold change (Log2)** | **P-Value(<0.05)** | **Protein description** |
| | **Iron ion transport** | | |
| Q920H8 | -3.90761 | 0.00053 | Hephaestin (Heph) |
| A0A0G2QC06 | -2.49118 | 0.00013 | Signal recognition particle receptor subunit beta (Tf) |
| A0A0G2K9I6 | -1.81357 | 0.00041 | Ceruloplasmin (Cp) |
| **Complement activation, classical pathway** | | | |
| D3ZFF8 | -4.67704 | 0.01268 | Ig-like domain-containing protein |
| D3ZWD6 | -3.66373 | 0.00089 | Complement C8 alpha chain (C8a) |
| A0A0G2K135 | -3.49305 | 0.00028 | Complement factor I (Cfi) |
| F1LZ11 | -3.49056 | 0.00186 | Ig-like domain-containing protein |
| A0A0G2K7X7 | -3.17118 | 0.00021 | Complement C7 (C7) |
| M0RBF1 | -3.05236 | 0.00050 | Complement C3 (C3) |
| P01026 | -2.58470 | 0.00106 | Complement C3 (C3) |
| Q62930 | -2.33416 | 0.00806 | Complement component C9 (C9) |
| Q6MG73 | -2.18306 | 0.00222 | C3/C5 convertase (C2) |
| A0A0G2K421 | -1.83630 | 0.00160 | Ig-like domain-containing protein |
| A0A0U1RRP9 | -1.43646 | 0.00092 | C3/C5 convertase (Cfb) |
| M0RB00 | -1.07335 | 0.00263 | Complement C4A (C4a) |
| **Response to nutrient levels** | | | |
| A0A0G2K845 | -3.45104 | 0.00015 | Adiponectin, C1Q and collagen domain-containing (Adipoq) |
| D3ZUQ8 | -2.43608 | 0.00060 | SZT2 subunit of KICSTOR complex (Szt2) |
| F7EY92 | -2.19651 | 0.00039 | Methyl-CpG binding domain protein 3 (Predicted), isoform CRA_c (Mbd3) |
| Q8R4Z9 | -1.74053 | 0.00071 | Mitofusin-1 (Mfn1) |
| F1LM16 | -0.92973 | 0.04693 | Plasminogen activator inhibitor 1 (Serpine1) |
| P01346 | -0.83931 | 0.01811 | Insulin-like growth factor II (Igf2) |
| P14841 | -0.61794 | 0.01409 | Cystatin-C (Cst3) |
| P04177 | -0.14992 | 0.04806 | Tyrosine 3-monooxygenase (Th) |
| **Response to organic cyclic compound** | | | |
| F1LNY6 | -4.43597 | 0.00095 | Transcription initiation factor TFIID 150 kDa subunit (Taf2) |
| P06866 | -4.01328 | 0.00008 | Haptoglobin (Hp) |
| Q80ZA3 | -3.07475 | 0.00328 | Alpha-2 antiplasmin (Serpinf1) |
| A0A0G2QC06 | -2.49118 | 0.00013 | Signal recognition particle receptor subunit beta (Tf) |
| F7EY92 | -2.19651 | 0.00039 | Methyl-CpG binding domain protein 3 (Predicted), isoform CRA_c (Mbd3) |
| P01346 | -0.83931 | 0.01811 | Insulin-like growth factor II (Igf2) |
| P14841 | -0.61794 | 0.01409 | Cystatin-C (Cst3) |
| P04177 | -0.14992 | 0.04806 | Tyrosine 3-monooxygenase (Th) |
| **Response to estradiol** | | | |
| O70249 | -3.63908 | 0.00023 | N-glycosylase/DNA lyase (Ogg1) |
| M0RBF1 | -3.05236 | 0.00050 | Complement C3 (C3) |
| P01026 | -2.58470 | 0.00106 | Complement C3 (C3) |
| P01346 | -0.83931 | 0.01811 | Insulin-like growth factor II (Igf2) |
| Q7TMA5 | -1.77291 | 0.00049 | Apolipoprotein B-100 (Apob) |
| A0A0G2K5J1 | -1.25927 | 0.00285 | Ryanodine receptor 1 (Ryr1) |
| P14841 | -0.61794 | 0.01409 | Cystatin-C (Cst3) |
| P04177 | -0.14992 | 0.04806 | Tyrosine 3-monooxygenase (Th) |
| F7EY92 | -2.19651 | 0.00039 | Methyl-CpG binding domain protein 3 (Predicted), isoform CRA_c (Mbd3) |
| **Protein transport** | | | |
| A0A0G2JU49 | 0.43709 | 0.00239 | Metaxin 1 (Mtx1) |
| D4A631 | 0.44235 | 0.00829 | Brefeldin A-inhibited guanine nucleotide-exchange protein 1 (Arfgef1) |
| A0A0G2JYT1 | 0.44303 | 0.00454 | ELKS/Rab6-interacting/CAST family member 1 (Erc1) |
| P11654 | 0.44750 | 0.02197 | Nuclear pore membrane glycoprotein 210 (Nup210) |
| Q6P756 | 0.44805 | 0.00339 | Adaptin ear-binding coat-associated protein 2 (Necap2) |
| D3ZT01 | 0.46375 | 0.00151 | Component of oligomeric Golgi complex 2 (Cog2) |
| Q5U316 | 0.46913 | 0.00176 | Ras-related protein Rab-35 (Rab35) |
| O35550 | 0.48829 | 0.00151 | Rab GTPase-binding effector protein 1 (Rabep1) |
| G3V870 | 0.50041 | 0.01281 | RCG51060 (Spire2) |
| Q66H90 | 0.53576 | 0.04901 | Bardet-Biedl syndrome 7 protein homolog (Bbs7) |
| Q6P6S4 | 0.55432 | 0.00107 | Nucleotide exchange factor SIL1 (Sil1) |
| Q8R491 | 0.56261 | 0.00781 | EH domain-containing protein 3 (Ehd3) |
| P29067 | 0.58863 | 0.00002 | Beta-arrestin-2 (Arrb2) |
| Q5U2U4 | 0.62999 | 0.00770 | Secretory carrier-associated membrane protein (Scamp2) |
| Q6PEC3 | 0.68624 | 0.00109 | Protein YIF1B (Yif1b) |
| P45479 | 0.75598 | 0.01011 | Palmitoyl-protein thioesterase 1 (Ppt1) |
| F1LR42 | 0.89870 | 0.00349 | RUN and FYVE domain-containing 1 (Rufy1) |

*Table S1. Proteins involved in multiple biological pathways were significantly changed in cells exposed to too low or pathologically high concentrations of tyrosine.*

*Table S2. Complete list of significantly altered protein abundances in response to either low or high environmental L-Tyr concentrations. See data in separate Excel file.*

**Modeling of TH activity and DA homeostasis**

We performed mathematical modeling to account for changes in TH activity and DA homeostasis in response to amino acid precursor manipulation. The different reactions were modelled using a differential equation approach. The rate equations were incorporated into a two-compartment model; cytosol and a vesicular compartment of the dopamine containing large dense-core vesicles, in the simulation software Copasi (v4.35, (Hoops et al., 2006)). Copasi uses the LSODA algorithm for stiff- and non-stiff systems and standard settings were used for numerical integration. The model presented here has implemented rate equations to describe TH activity at conditions where BH4, Tyr, Phe levels vary in addition to accounting for varying feedback inhibition by dopamine (DA) and different levels of Ser40 phosphorylation (see Table S3 for parameter values).

Our model differ from previous models of dopamine homeostasis (Best et al., 2010, 2009) for several reasons. The models from Best et al. are addressing striatal dopamine, whereas we have used PC12 cells in this study. In the Best model, Tyr was partially unavailable for dopamine synthesis by separation into a “storage pool”, which is not well described experimentally and can complicate the behavior of the model, especially when the effect of cellular amino acid levels is to be modelled. We therefore treated all substrates as well mixed and available to their enzymes within the compartments included. The only exception to this was the vesicular DA, which was largely associated with a vesicular matrix to allow the high storage concentrations reported in the literature. This is in line with experimental findings on synaptic and dense-core vesicles in chromaffin and in PC12 cells.

The modeling of vesicular storage in Best et al. gives only 81 $\mu$M DA within the vesicles, which is about three orders of magnitude lower than experimental measurements: 60-100 mM in PC12 cells (Li et al., 2018) and about 100 mM in striatal dopamine terminal synaptic vesicles (Omiatek et al., 2013). The typical size of the PC12 large dense-core vesicles where DA is stored is 160-215 nm (diameter) (Dembla & Becherer, 2021), giving a volume of about 4.2 al/vesicle. A total pool of large dense-core vesicles is estimated to about 7000/cell for PC12, with a population of about 1000/cell in the readily realizable pool (Kasai et al., 1996). This gives a total volume of 29.4 fl for the vesicular compartment. The total volume of a typical PC12 cell (10-15 $\mu$m) is about 1.02 pl, where about 20 % of the volume is assumed to be contained by the nucleus and other organelles. The cytosolic volume of 817.6 fl is then 96.5 % of the total two-compartment volume and the large dense-core vesicular volume is 3.5 % of the total model volume. Under the assumption that the concentration of DA is about 1.0 $\mu$M in the cytosol and 80 mM in the large dense-core vesicles, the cytosol contain only 0.035 % of the total cellular DA and 14.3 % of the total DA is in the readily realizable vesicular pool. To achieve a concentration of about 80 mM within the vesicles, DA is co-stored with a matrix to increase the solubility in an acidic environment where DA is stable. We have modelled this as the formation of a DA-matrix complex within the vesicles with rate constants that does not limit other processes (see Table S4 and text).

*Table S3: Modeling of TH activity and kinetics*

The table summarizes kinetic parameters for TH used in the modeling and literature values reported with their corresponding reference. The values in bold are used in the model. The reactions involving

TH used in the model are: Phe + BH4 -> Tyr + 4aOH-BH4 for TH-mediated phenylalanine hydroxylation (rate: $v_{TH,Tyr}$) and Tyr + BH4 -> Dopa + 4aOH-BH4 for tyrosine hydroxylation (rate: $v_{TH,Dopa}$). BH4 regeneration is included in the model by a 4aOH-BH4 -> BH4 reaction (v = 3.33 $min^{-1}$ [4aOH-BH4]; where the rate constant is set to the $\boldsymbol{V_{max}^{f}}$ value of Dihydropteridine reductase in Best et al. 2009)). The $K_m$ values for different substrates (S) are given as $\boldsymbol{K_m^S}$, whereas $\boldsymbol{K_{si}^{Tyr}}$ is the substrate inhibition constant for Tyr, and $\boldsymbol{K_{i,(p)TH}^{DA}}$ is the inhibition constants for DA. Kinetic values that differ between non-phosphorylated TH and Ser40 phosphorylated TH are labelled with subscript TH or pTH, respectively. $V_{max}$ values are provided as (%) relative to that of non-phosphorylated TH with Tyr as substrate. The $V_{max}$ value for non-phosphorylated TH is set to 2.08 μM/min in the model (same as 125 μM/hr used in Best et al. 2009). In the model where spontaneous release of vesicular DA is included, the $V_{max}$ value for TH was increased 100-fold to 208 μM/min. All other parameters were unchanged. The fraction of TH which is Ser40 phosphorylated is given by $r_p$.

| **Parameters** | **Values** | **References** |
|---|---|---|
| $\boldsymbol{K_m^{Tyr}}$ (μM) | 16; 51; 5.1; 17; 46; 31<br>Av. 27; Median **24** | (Barone et al., 2020; Calvo et al., 2010; Daubner et al., 2000; Daubner & Fitzpatrick, 1998; Fossbakk et al., 2014; Royo et al., 2005) |
| $\boldsymbol{K_m^{Phe}}$ (μM) | 100; 109; 296; 100<br>Av. 151; Median **104** | (Daubner et al., 2000; Daubner & Fitzpatrick, 1998; Fossbakk et al., 2014; Ribeiro et al., 1991) |
| $\boldsymbol{K_{m,TH}^{BH4}}$ (μM) | 39; 46; 13; 27; 24; 24<br>Av. 28; Median **26** | (Calvo et al., 2010; Daubner & Fitzpatrick, 1998; Flatmark et al., 1999; Fossbakk et al., 2014; Royo et al., 2005; Toska et al., 2002) |
| $\boldsymbol{K_{m,pTH}^{BH4}}$ (μM) | **8** | (Toska et al., 2002) |
| $\boldsymbol{K_{si}^{Tyr}}$ (μM) | 59; 92; 46; 29<br>Av. 57; Median **53** | (Barone et al., 2020; Calvo et al., 2010; Fossbakk et al., 2014; Royo et al., 2005) |
| $\boldsymbol{K_{i,TH}^{DA}}$ (nM) | 56.4*; 0.71$<br>**6.3¶** | (Bueno-Carrasco et al., 2022; Ramsey & Fitzpatrick, 1998) |
| $\boldsymbol{K_{i,pTH}^{DA}}$ (nM) | 477*; 220$<br>**324¶** | (Bueno-Carrasco et al., 2022; Ramsey & Fitzpatrick, 1998) |
| $\boldsymbol{V_{max,pTH}^{Tyr}}$(% of $\boldsymbol{V_{max,TH}^{Tyr}}$) | **160** | (Toska et al., 2002) |
| $\boldsymbol{V_{max}^{Phe}}$ (% of Tyr) | 64; 64; 67<br>Av. **65** | (Daubner et al., 2000; Daubner & Fitzpatrick, 1998; Fossbakk et al., 2014) |
| $\boldsymbol{r_p}$ | 0.05 basal<br>0.15 stimulated | (Dunkley et al., 2004) |

*calculated from reported $IC_{50}$ values of 0.49 and 12.4 μM, using the relation $K_i = IC_{50}*K_m/(K_m+[S])$, where [BH4] = 200 μM, $K_m$ = 26 μM for TH and 8 μM for pTH. $reported as $K_d$ values. ¶calculated as geometric mean of the two literature values.

*Rate equations used for the modeling*

$$v_{TH,Dopa} = \left( (1-r_p)V_{max,TH}^{Tyr} \left( \frac{BH4}{K_{m,TH}^{BH4}\left(1+\left(\frac{DA}{K_{i,TH}^{DA}}\right)\right)+BH4} \right) + r_p V_{max,pTH}^{Tyr} \left( \frac{BH4}{K_{m,pTH}^{BH4}\left(1+\left(\frac{DA}{K_{i,pTH}^{DA}}\right)\right)+BH4} \right) \right) \left( \frac{Tyr}{K_m^{Tyr}\left(1+\left(\frac{Phe}{K_{i2}^{Phe}}\right)\right)+Tyr\left(1+\left(\frac{Tyr}{K_{si}^{Tyr}}\right)\right)} \right) \quad (1)$$

$$v_{TH,Tyr} = \left( (1-r_p)V_{max,TH}^{Phe} \left( \frac{BH4}{K_{m,TH}^{BH4}\left(1+\left(\frac{DA}{K_{i,TH}^{DA}}\right)\right)+BH4} \right) + r_p V_{max,pTH}^{Phe} \left( \frac{BH4}{K_{m,pTH}^{BH4}\left(1+\left(\frac{DA}{K_{i,pTH}^{DA}}\right)\right)+BH4} \right) \right) \left( \frac{Phe}{K_m^{Phe}\left(1+\left(\frac{Tyr}{K_i^{Tyr}}\right)\right)+Phe} \right) \quad (2)$$

$$v_{DDC} = V_{max}^{DDC}\left(\frac{Dopa}{K_m^{DDC}+Dopa}\right) \quad (3)$$

$$v_{VMAT2} = V_{max}^{VMAT2}\left(\frac{DA}{K_m^{VMAT2}+DA}\right) \quad (4)$$

$$v_{ves.leak} = k_{leak}^{ves} DA_v \quad (5)$$

$$v_{MAO} = k_{met}^{MAO} DA \quad (6)$$

$$v_{DArel} = k_{rel}[DA_v{:}M_v] \quad (7)$$

*Table S4. Model reactions and parameter values not involving TH*

The concentration of DA binding sites in the vesicular matrix ($M_v$) were set to 80 mM. The parameter values in parentheses represent the values used in the model corrected for spontaneous release of vesicular DA (10-fold higher than in the original model).

| Enzyme/process | Reactions | Parameters | Values | References |
|---|---|---|---|---|
| DDC | Dopa -> DA | $K_m^{DDC}$ (μM) | 130 | (Best 2009) |
| | | $V_{max}^{DDC}$ (μM/min) | 167 (1670) | (Best 2009) This study |
| VMAT2 | DA -> $DA_v$ | $K_m^{VMAT2}$ (μM) | 3.00 | (Best 2009) |
| | | $V_{max}^{VMAT2}$ (μM/min) | 118 (1180) | (Best 2009) This study |
| Leakage of unbound vesicular DA to the cytosol | $DA_v$ -> DA | $k_{leak}^{ves}$ | 0.667 (6.67) | (Best 2009) This study |
| Binding and unbinding of DA to vesicular matrix | $DA_v$ + $M_v$ -> $DA_v$:$M_v$ | $k_a^{Mv}$ (μM$^{-1}$min$^{-1}$) | 100 | This study |
| | $DA_v$:$M_v$ -> $DA_v$ + $M_v$ | $k_d^{Mv}$ (min$^{-1}$) | 10.0 | This study |
| Metabolism and oxidation of cytosolic DA | DA -> | $k_{met}^{MAO}$ | 8.69 $10^{-2}$ (0.869) | This study This study |
| *Spontaneous release of* vesicular DA* | *$DA_v$:$M_v$ -> $M_v$* | *$k_{rel}$ (min$^{-1}$)* | *0.007* | *See text* |

*This process was only included in the model with high $V_{max}$ values and rate constants in ().

| Metabolite/flux | Value |
|---|---|
| Phe (μM) | 0 |
| Tyr (μM) | 126 |
| BH4 (μM) | 359.97 |
| 4aOH-BH4 (nM) | 26.1 |
| Dopa (nM) | 83.3 |
| DA (μM) | 1.00 |
| $DA_v$ (μM) | 44.2 |
| $DA_v$:$M_v$ (mM) | 79.8 |
| $M_v$ (mM) | 0.180 |
| $v_{TH,Dopa}$ (μM/min) | 8.69 $10^{-2}$ |

*Table S5. Steady state metabolite concentrations and TH activity for the initial model at the Tyr and BH4 levels used by Best et al. 2009*

Using the model setup described we set the [Phe] = 0, [Tyr] = 126 μM and [BH4] = 360 μM (concentrations used in the model by Best et al. (Best et al., 2009) to adjust the rate constant $k_{met}^{MAO}$ of cytosolic DA metabolism (and oxidation) so that the cytosolic DA steady state level reached 1.00 μM in absence of any release of DA and uptake of extracellular DA (Table S5). This steady state level of cytosolic DA is lower than in the Best model (2.65 μM), but forcing it to this level in our model, would reduce $k_{met}^{MAO}$ to 2.15 $10^{-2}$ min$^{-1}$. At this condition, the flux through TH Dopa synthesis was 5.2 μM/hr compared to 27 μM/hr in the Best model (Table S5). This is logical as a stronger inhibition by DA is included in our model as well as a stronger substrate inhibition by tyrosine. A lower DA synthesis flux is also expected in PC12 compared to striatal DA terminals and in absence of DA release.

The PC12 cell has a reported spontaneous secretion rate of about 12.7 pmol/$10^5$ cells in 10 min and about 50 pmol/$10^5$ cells during 56 mM KCl stimulation (Ritchie, 1979). This corresponds roughly to 50 and 200 vesicles min$^{-1}$ for spontaneous versus KCl stimulated conditions, or about 0.7 and 3 % of the total pool (5 or 20 % or the readily releasable pool) of vesicles min$^{-1}$, respectively. This corresponds to 10 times turnover of the vesicular pool during 24 h for the spontaneous release rate. We added release of DA to the model by the following reaction: $DA_v$:$M_v$ -> $M_v$ where the rate is first order kinetics. In addition, the PC12 cells grow quite fast, expanding at least two-fold every 24 h. We assume, however, that the proteins involved are expressed at the same level during cell growth and therefore the $V_{max}$ values are scaled accordingly and stay constant during cell division. Still, this would mean a doubling of the vesicular pool every 24 h with vesicles that needs filling of DA. This rate is however small compared to the turnover of vesicular DA through spontaneous release and was therefore ignored here.

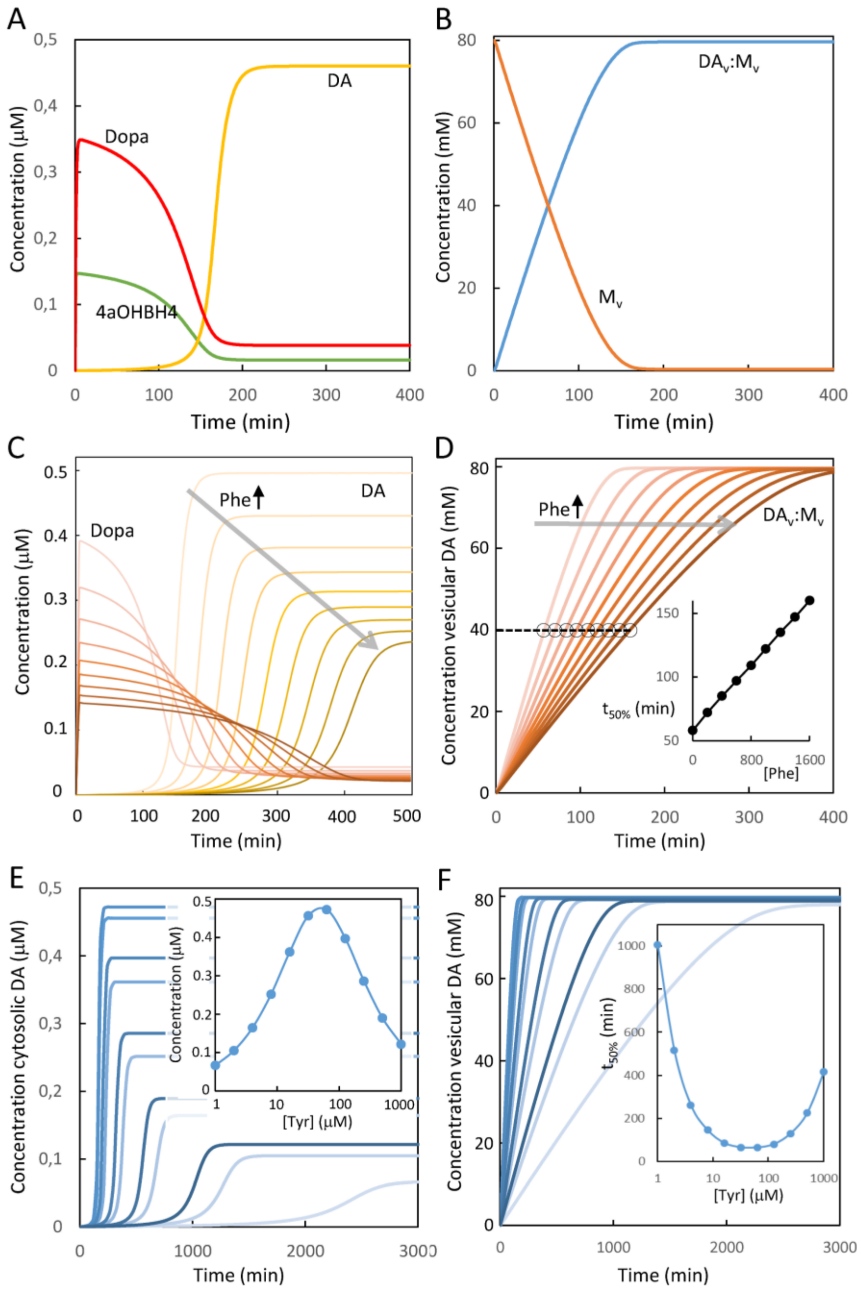


*Fig. S5. Behavior of the initial model of dopamine homeostasis. The model was run to steady state after emptying a smaller part (1.5 %) of the total vesicular pool, starting from an initial state (intracellular Phe and Tyr fixed at 100 and 75 µM, respectively) without Dopa, cytosolic DA or vesicular DA. (A) shows the temporal change in Dopa (red), 4a-OH-BH4 (green) and cytosolic DA (yellow). (B) shows the temporal change in vesicular DA ($DA_v$) associating to the matrix ($M_v$, orange), forming vesicular matrix-bound DA ($DA_v$:$M_v$, blue). The majority of vesicular DA were in the form of matrix-bound DA (DAv:Mv, 79.7 mM), whereas unbound vesicular DA remained low in comparison (23.5 µM). In (C-F) the situations were modelled where cells were simultaneously challenged with a sudden change in cellular Phe (C and D) 0-1600 µM or Tyr (E and F) 1-1000 µM, color coded from light (low Phe/Tyr) to dark (high Phe/Tyr). (C) shows the temporal change in Dopa (brown) or cytosolic DA (yellow) upon gaining steady state at different cytosolic conc. of Phe. (D) shows the simultaneous change in $DA_v$:$M_v$, with the inset showing the time needed for 50% filling of $DA_v$:$M_v$ for different cytosolic conc. of Phe. The concentration in vesicular DA is followed over time (color coded from low (light) to high (dark) conc.). Panels (E) and (F) show the steady state conc. of vesicular DA at different cellular conc. of Tyr.*

**Modeling results**

*Initial modeling without continuous vesicular DA release*

We modelled first the situation where the cells were emptied of 1.5 % its total vesicular pool or about 10 % of the readily releasable vesicular pool, without any further release of DA and without reuptake of extracellular DA. The simulation was started without any cytosolic DA or Dopa, thus relying only on new biosynthesis and the basal physiological state was set to 26 μM BH4 (equal to the $K_m$ value), 100 μM Phe and 75 μM Tyr. The model predicted a cytosolic DA steady state level of 0.46 μM (Table S6, Fig. S5A,B). During the pre-steady state phase, the emptied vesicular pool recruited newly synthesized DA, forming matrix-associated DA and this prevented cytosolic accumulation and feedback inhibition of TH. Half refilling was achieved after approximately 63 min at our physiological modeling conditions (Fig. S5B).

We further modelled the situation where the cytosolic levels of Tyr and Phe were clamped at different concentrations and then monitored how the steady state level of the other metabolites of the DA pathway changed as well as the flux of TH-mediated Tyr and Dopa synthesis (Table S6, Fig. S5C-F). Increasing the cellular Phe level 0-1600 μM (with Tyr clamped at 75 μM) slowed down the kinetics of vesicular DA filling (Fig. S5D), with a linear increase in time needed for 50 % vesicular filling versus the cellular concentration of Phe (Inset, Fig. S5D). The steady state cytosolic DA level decreased also with increasing Phe from about 0.5 μM to less than 0.25 μM (Fig. S5C). Tyrosine had a larger and biphasic impact on both steady state cytosolic DA levels (Fig. S5E and inset) and the time needed for 50 % vesicular refilling (Fig. S5F and inset). Of note, the time of 50 % vesicular refilling for different Tyr concentrations was rather flat at the bottom with little difference between 10-200 μM Tyr, but with larger effects at very low and very high Tyr levels (Inset, Fig. S5F). This shape was different to the response curve for cytosolic DA to Tyr concentration (Inset, Fig. S5E). The optimal refilling rate corresponded well to the substrate inhibition behavior of TH, suggesting that this feature could well impact the cellular capacity to replenish its vesicular DA pool. We noted, however, that the initial model without continuous release of vesicular DA did not predict any significant change in steady state vesicular DA and therefore no significant change in total cellular DA in response to changes in Tyr or Phe levels. Still, as the rate of refilling was very slow, which could explain experimental measurements performed here as being a result of steady state and pre-steady state measurements. We find this to be rather unlikely, though.

*Table S6. Steady state concentrations and flux at different conditions*

The table shows the output of the model for different clamped levels of Phe and Tyr. Condition 0 corresponds to the conditions used in Best et al. 2009 for Phe, Tyr and BH4 and the computed steady state metabolite and fluxes in our model. Conditions 1-4 are computed values for physiological and high levels of Phe or Tyr. For condition 5 the cellular BH4 level is doubled and in condition 6 the Ser40 phosphorylation of TH is increased 4-fold relative to basal. The modeling performed here was performed in absence of secretion of vesicular or uptake of extracellular DA.

| **Metabolite/flux** | **Cond. 1** | **Cond. 2** | **Cond. 3** | **Cond. 4** | **Cond. 5** | **Cond. 6** |
|---|---|---|---|---|---|---|
| Phe (μM)* | 100 | 1000 | 100 | 1000 | 100 | 100 |
| Tyr (μM)* | 75 | 75 | 835 | 835 | 75 | 75 |
| BH4 (μM) | 25.98 | 25.99 | 25.99 | 25.99 | 51.98 | 25.96 |
| 4aOH-BH4 (nM) | 16.1 | 12.1 | 4.65 | 7.82 | 20.7 | 38.7 |
| Dopa (nM) | 38.3 | 24.1 | 11.5 | 11.5 | 49.4 | 91.9 |
| DA (μM) | 0.460 | 0.290 | 0.131 | 0.127 | 0.59 | 1.10 |
| $DA_v$ (μM) | 25.5 | 15.6 | 7.40 | 7.21 | 29.2 | 47.58 |
| $DA_v:M_v$ (mM) | 79.7 | 79.5 | 78.9 | 78.9 | 79.7 | 79.8 |
| $M_v$ (mM) | 0.338 | 0.510 | 1.07 | 1.10 | 0.273 | 0.168 |
| $v_{TH,Dopa}$ (μM/min) | $4.00\ 10^{-2}$ | $2.52\ 10^{-2}$ | $1.21\ 10^{-2}$ | $1.20\ 10^{-2}$ | $5.15\ 10^{-2}$ | $9.59\ 10^{-2}$ |
| $v_{TH,Tyr}$ (μM/min) | $1.37\ 10^{-2}$ | $1.52\ 10^{-2}$ | $3.40\ 10^{-3}$ | $1.40\ 10^{-2}$ | $1.76\ 10^{-2}$ | $3.28\ 10^{-2}$ |

*values fixed in the model (cell clamp).

*Modeling with spontaneous and continuous release of vesicular DA*

The initial model, emptying of 1.5 % of the total vesicular pool corresponds to about 105 vesicles. For that model operating at physiological conditions, a rate of replenishment of 52.5 vesicles in 63 min was estimated. This corresponds to a rate of refilling of only 50 vesicles/h. This is 60-fold slower than the rate of spontaneous release of vesicular DA, 50 vesicles/min, calculated based on literature values on the PC12 cell (methods/supplemental text). We therefore scaled up the $V_{max}$ values and rate constants of the initial model 100-fold for TH (see Table S3) and 10-fold for the other proteins (see Table S4). The $K_m$, $K_i$ and $K_{si}$ values were unchanged. See the results section of the main article for the remaining text on this model.

**Extending the model with amino acid uptake**

Aberrancies in amino acid transport into the brain and into monoamine producing cells has for long been viewed as one of the contributing mechanisms for neuropsychiatric comorbidities of PKU and HT-1. For the aromatic amino acids, LAT1 (SLC7A5) and LAT2 (SLC7A8) are key transporters, and the amino acids compete with transport into the cells. However, amino acid transporters generally have rather broad substrate specificity and transport several different amino acids. Therefore, the amino acid composition of the medium will affect the competitiveness between the aromatic amino acids. In fact, a quantitative description of amino acid homeostasis in cells is rather complex, but has been demonstrated to be feasible, provided a detailed knowledge of the expression levels of different amino acid transporters and their kinetics is known (Gauthier-Coles et al., 2021). We do not have a detailed expression profile for amino acid transporters in PC12 cells, so we have used the data of Gauthier-Coles et al. and experimental data on Tyr and Phe in PC12 cells from DePietro and Fernstrom (DePietro & Fernstrom, 1999) to derive a quantitative relation between extracellular and intracellular Phe and Tyr conc. at our growth conditions.

The data from DePietro and Fernstrom on PC12 cells suggest little impact of Tyr on the cellular Phe levels. Their data were used to estimate a relation between extracellular and intracellular Phe concentration (Fig. S6A). To derive this, we used the cell number to mg protein conversion of Gauthiers-Coles et al. (7 mill cells = 1 mg protein) as A549 and U87-MG cells have similar volume to PC12 cells (volume of 7 mill PC12 cells = 7.14 μl). Fitting the data with a hyperbolic function yielded a maximal intracellular Phe conc. of 1956 μM and a half max conc. at 260 μM extracellular Phe. This gives about 5.4-fold $Phe_{in}/Phe_{ex}$ at 100 μM $Phe_{ex}$ and 4.2-fold at 214 μM $Phe_{ex}$, which is close to 5.0 and 4.7-fold reported in U87-MG and A549 cells, respectively (Gauthier-Coles et al., 2021).

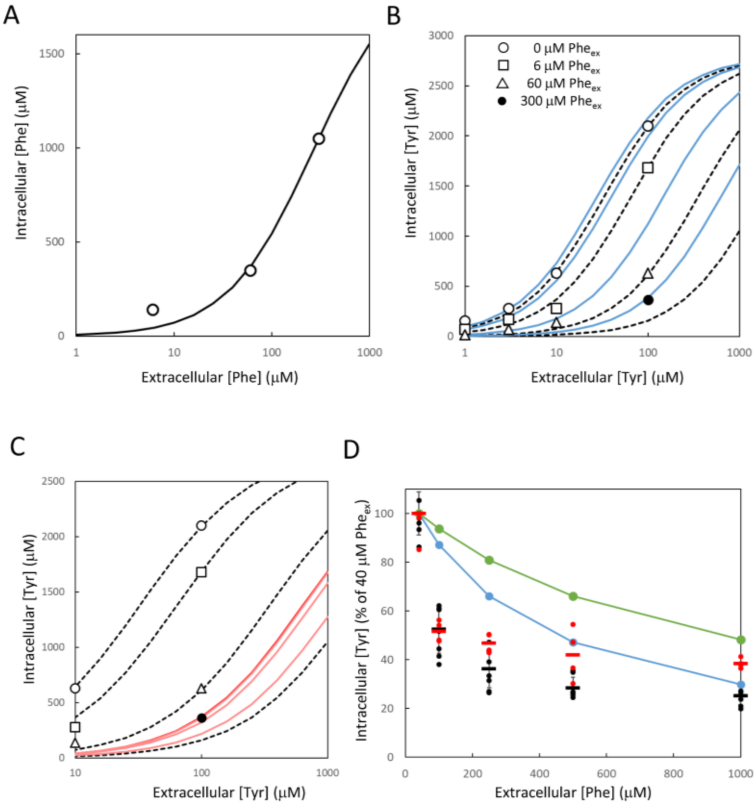


*Fig. S6. Modeling cellular Phe and Tyr transport. (A) shows the fitted model (black line) of intracellular Phe ($Phe_{in}$) as a function of extracellular Phe ($Phe_{ex}$) to experimental data (○) from DePietro and Fernstrom. (B) shows the best fitted model (dotted lines) of Tyr uptake for different conc. of $Phe_{ex}$ (data from DePietro and Fernstrom). The blue lines show a model where reported $K_m$ values for Phe and Tyr (Table S7) are used instead of the values obtained by best fit. (C) shows the predicted transport kinetics of Tyr (blue model in (B)) at different conc. of $Phe_{ex}$ (red lines, 0 μM $Phe_{ex}$ red, 6, 60, 300 μM $Phe_{ex}$ light red) in the presence of other amino acids at the conc. used here for PC12 cells (see Table S7). (D) shows the $Tyr_{in}$ (% of $Tyr_{in}$ at 40 μM $Phe_{ex}$) calculated for LAT1 (blue) or LAT2 (green) mediated transport, at different conc. $Phe_{ex}$ at the medium conditions used here (Table S7). Experimentally measured $Tyr_{in}$ levels (from Fig. 1C) are shown as black (1 and 3 h) and red (6 h) dots, with average values shown as (–).*

In their paper, DePietro and Fernstrom also measured $Tyr_{in}$ at different $Tyr_{ex}$ and $Phe_{ex}$ conc. These data could be fitted using competitive Michaelis Menten kinetics, giving an apparent $K_m$ value of 33.3 for $Tyr_{ex}$ and apparent $K_i$ of 6.2 for $Phe_{ex}$ (Fig. S6B, dotted lines). These values are quite close to the reported $K_m$ values of Tyr and Phe for LAT1 – 28 and 14 μM, respectively. Using these values gives a poorer, but still reasonable fit to the data, particularly for high Phe (Fig. S6B, blue lines). However, the experiments in DePietro and Fernstrom were performed in media depleted for any other amino acids. We therefore used the kinetic approach from Gauthier-Coles et al. to estimate the apparent $K_m$ values for Phe and Tyr for LAT1 and 2 at our growth conditions (Table S7, eq 8). For LAT1, His and Leu contributed the most to increase the apparent $K_m$ value, whereas Gln and Cys contributed most for LAT2.

*Table S7. Amino acid content of media and $K_m$ values for LAT1 and 2.*

| | Conc. in medium $[AA_i]$ (μM) | SLC7A5 LAT1 Km (μM) | SLC7A8 LAT2 $K_m$ (μM) | SLC7A5 LAT1 $[AA_i]/K_{mi}$ | SLC7A8 LAT2 $[AA_i]/K_{mi}$ |
|---|---|---|---|---|---|
| His | 124 | 12 | 181 | *10.3* | 0.685 |
| Ile | 52 | 25 | 97 | 2.08 | 0.536 |
| Leu | 131 | 20 | 119 | *6.55* | 1.10 |
| Met | 37 | 20 | 204 | 1.85 | 0.181 |
| Trp | 13.6 | 21 | 58 | 0.648 | 0.234 |
| Val | 58 | 47 | 124 | 1.23 | 0.468 |
| Ala | 163 | | 187 | | 0.872 |
| Asn | 103 | | 81 | | 1.27 |
| Cys | 218 | | 109 | | *2.00* |
| Gln | 1000 | | 151 | | *6.62* |
| Ser | 116 | | 116 | | 1.00 |
| Thr | 62 | | 69 | | 0.898 |
| Σ $[AAi]/K_{mi}$ | | | | 22.7 | 15.9 |
| **Phe** | **40** | **14** | **45** | **2.86** | **0.889** |
| **Tyr** | **35** | **28** | **36** | **1.25** | **0.972** |

$$K_{m,app}^{Tyr} = K_m^{Tyr}\left(1 + \frac{Phe_{ex}}{K_m^{Phe}} + \sum\frac{AA_i}{K_m^{AAi}}\right) \quad (8)$$

$$v_{LAT}^{Tyr} = V_{max}^{LAT}\frac{Tyr_{ex}}{K_{m,app}^{Tyr}+Tyr_{ex}} \quad (9)$$

$$v_{in}^{Tyr} = v_{LAT1}^{Tyr} + v_{LAT2}^{Tyr} \quad (10)$$

$$v_{LAT1,out}^{Tyr} = V_{max}^{LAT1}\frac{Tyr_{in}}{1000\,K_{m,app}^{LAT1}}, v_{LAT2,out}^{Tyr} = V_{max}^{LAT2}\frac{Tyr_{in}}{200\,K_{m,app}^{LAT2}} \quad (11)$$

$$v_{out}^{Tyr} = v_{LAT1,out}^{Tyr} + v_{LAT2,out}^{Tyr} + V_{max}^{out}\frac{Tyr_{in}}{K_{m,app,out}^{Tyr}+Tyr_{in}} \quad (12)$$

Using this approach, we calculated the impact of the other amino acids present in the growth medium assuming that only LAT1 (80 % of $V_{max}$) and LAT2 (20 % of $V_{max}$) contributed to the inward transport (Eq. 10). The predicted response curves at different $Phe_{ex}$ conc. becomes very right-shifted (high apparent $K_m$ values) from the fitted curves in absence of other amino acids (dotted lines) and much less responsive to $Phe_{ex}$, with 60 μM Phe very slightly right-shifted and 300 μM more so (Fig. S6C). This means that overlapping substrate specificity of the transporters is making the cellular amino acid homeostasis more robust towards changes in the level of a single amino acid. Also, the growth conditions and cell model would make an impact on the intracellular Tyr levels and therefore the expected impact on the biosynthesis. We also calculated the % change in LAT1 and LAT2 Tyr-transport activity (relative to that at 40 μM $Phe_{ex}$) at different $Phe_{ex}$ conc. for the growth conditions used here (Fig. S6D, blue and green, respectively). The experimental differences in cellular Tyr levels (from Fig. 1C) showed a steeper response to $Phe_{ex}$ than the stipulated LAT1 and 2 activities. We therefore added outward Tyr transport comprised of LAT1 and 2 activity (reversible transporters) and an additional transporter with Michaelis Menten kinetics (Eq. 11 and 12). The amino acid transport models for Phe and Tyr were used to calculate $Phe_{in}$ and $Tyr_{in}$ values for input to the DA homeostasis model. Three models with different parameter sets of $V_{max}^{out}$ and $K_{m,app,out}^{Tyr}$ were generated based on their fit to the data on DA levels (data from Fig. 1A and B) at different $Phe_{ex}$ (Fig. S7A, 7A) and $Tyr_{ex}$ (Fig. S7B, 7B) levels. See the results section of the paper for further details.

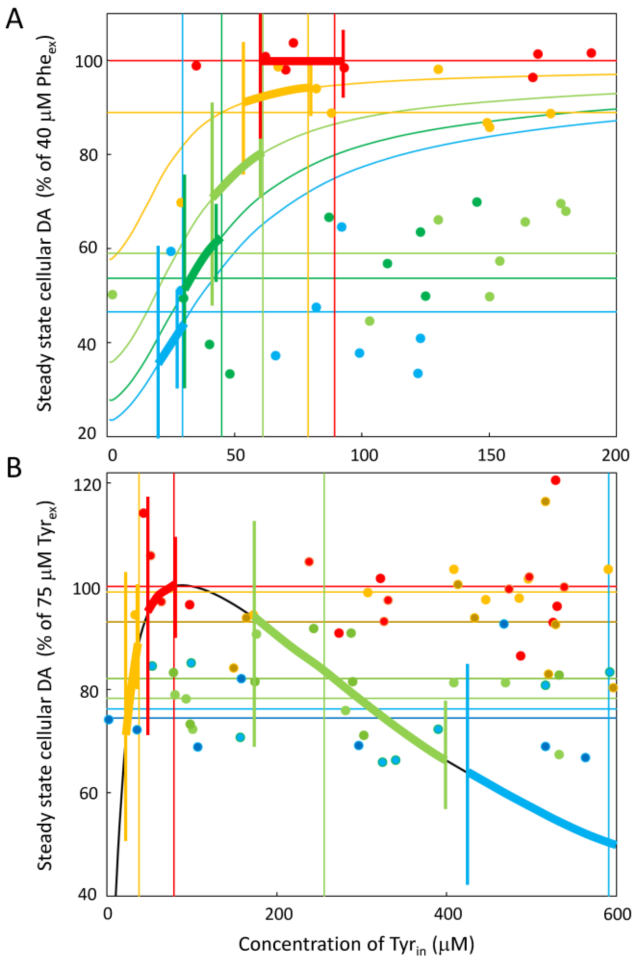


*Fig. S7. Comparison between experimental data and model predictions. The steady state cellular DA levels (% of (A) that for 40 μM $Phe_{ex}$ and (B) that of 75 μM $Tyr_{ex}$) is shown for (A) different conc. of $Phe_{ex}$ ($Tyr_{ex}$ 75 μM) and (B) different conc. of $Tyr_{in}$ ($Phe_{ex}$ 100 μM). Experimental data is shown as dots, colored by different conditions and with the average values shown as horizontal lines. The predicted values of cellular DA from the DA synthesis model are shown as lines for varying levels of $Tyr_{in}$. In (A) the calculations were performed for the different predicted $Phe_{in}$ values for $Phe_{ex}$ values of 40 μM (261 μM $Phe_{in}$, red = 100 %), 100 μM ($Phe_{in}$ 543.8 μM, orange), 250 μM ($Phe_{in}$ 959.4 μM, light green), 500 μM ($Phe_{in}$ 1287.4 μM, green) and 1000 μM ($Phe_{in}$ 1552.8 μM, blue). The same color codes are used for the experimental data (dots, average as horizontal lines). (B) shows the data for $Tyr_{ex}$ 35 μM (orange), 75 μM (red, 100 %), 275 μM (green) and 835 μM (blue). The model prediction for varying $Tyr_{in}$ levels ($Phe_{ex}$ 100 μM/ $Phe_{in}$ 543.8 μM) is shown as a black line. The $Tyr_{in}$ values predicted by the three different transport models for each condition (in (A) varying $Phe_{ex}$ and (B) varying $Tyr_{ex}$) is shown as vertical lines, for different parameter set of $V_{max}^{out}$/ $K_{m,app,out}^{Tyr}$ (short lines, 50 % of $V_{max}^{in}$ /400 μM; full vertical lines, 180 % of $V_{max}^{in}$ /1100 μM; medium length lines, 220 % of $V_{max}^{in}$ /2000 μM). The area of predicted DA levels for the three transport models is between the points where the vertical lines cross their corresponding model prediction line (shown as fat lines). The range of predicted DA levels can then be compared to the range of values obtained experimentally (colored dots). In (B), the predicted $Tyr_{in}$ for the low $V_{max}^{out}$/ $K_{m,app,out}^{Tyr}$ model for $Tyr_{ex}$ of 835 μM was 9631 μM, which has a predicted % DA level of 2.52, and is therefore outside the scale of the figure.*
